# Architecture and self-assembly of the jumbo bacteriophage nuclear shell

**DOI:** 10.1101/2022.02.14.480162

**Authors:** Thomas G. Laughlin, Amar Deep, Amy M. Prichard, Christian Seitz, Yajie Gu, Eray Enustun, Sergey Suslov, Kanika Khanna, Erica A. Birkholz, Emily Armbruster, J. Andrew McCammon, Rommie E. Amaro, Joe Pogliano, Kevin D. Corbett, Elizabeth Villa

**Affiliations:** Division of Biological Sciences, University of California San Diego, La Jolla, CA, USA; Department of Cellular and Molecular Medicine, University of California San Diego, La Jolla, California, USA; Department of Chemistry and Biochemistry, University of California San Diego, La Jolla, California, USA; Department of Pharmacology, University of California San Diego, La Jolla, California, USA; Howard Hughes Medical Institute, University of California San Diego, La Jolla, California, USA

## Abstract

Bacteria encode myriad defenses that target the genomes of infecting bacteriophage, including restriction-modification and CRISPR/Cas systems. In response, one family of large bacteriophage employs a nucleus-like compartment to protect their replicating genomes by excluding host defense factors. However, the principle composition and structure of this compartment remain unknown. Here, we find that the bacteriophage nuclear shell assembles primarily from one protein, termed chimallin. Combining cryo-electron tomography of nuclear shells in bacteriophage-infected cells and cryo-electron microscopy of a minimal chimallin compartment in vitro, we show that chimallin cooperatively self-assembles as a flexible sheet into closed micron-scale compartments. The architecture and assembly dynamics of the chimallin shell suggest mechanisms for its nucleation and growth, and its role as a scaffold for phage-encoded factors mediating macromolecular transport, cytoskeletal interactions, and viral maturation.

## Introduction

Over billions of years of conflict with bacteriophages (phages), plasmids, and other mobile genetic elements, bacteria have evolved an array of defensive systems to target and destroy foreign nucleic acids (1). Phages have in turn evolved mechanisms, including anti-restriction and anti-CRISPR proteins, that counter specific bacterial defense systems (5–7). We recently showed that a family of “jumbo phages” – named for their large genomes (typically >200 kb) and large virion size – assemble a selectively-permeable, protein-based shell that encloses the replicating viral genome and is associated with a unique phage life cycle, represented in (**Fig. 1a**) (2, 8).

**Figure 1.**
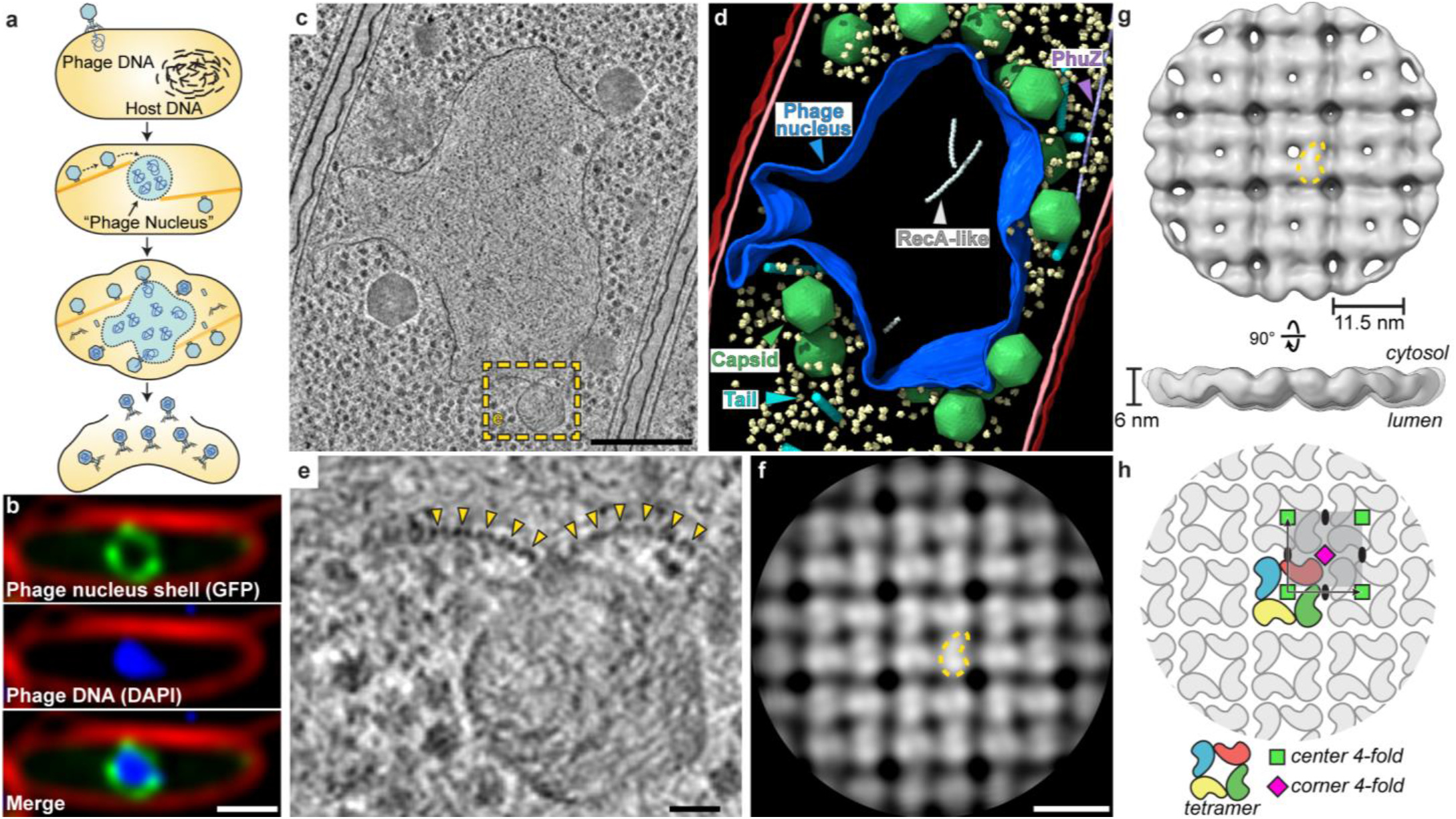
*In situ* tomography and subtomogram analysis of the 201φ2-1 phage nucleus. **a**, Schematic of the jumbo phage infection cycle. **b**, Fluorescence microscopy of a 201φ2-1-infected *P. chlororaphis* cell at 45 mpi. Phage nucleus shell component, gp105, (green) is tagged with GFP, phage DNA (blue) is stained with DAPI, and the outer cell membrane (red) is stained with FM4-64. **c**, Tomographic slice of a phage nucleus in a 201φ2-1-infected *P. chlororaphis* cell. **d**, Segmentation of the tomogram in **c**. Outer and inner bacterial membranes are burgundy and pink, respectively. The phage nucleus is colored blue. Phage capsids and tails are green and cyan, respectively. PhuZ and recA-like protein filaments are light purple and white, respectively. A subset of five-hundred host ribosomes are shown in pale-yellow. **e**, Enlarged view of the boxed region in **c**. Yellow arrows point to the repetitive feature of the phage nucleus perimeter. **f**, Slice of the cytosolic face of the subtomogram average of the repetitive feature in the phage nucleus perimeter with a comma-shaped subunit outlined in yellow. **g**, Cytosolic and side views of the shell subtomogram average isosurface with a single subunit outlined in yellow. **h**, Schematic representation of the *p442*-like arrangement of chimallin protomers. Scale bars: **b**:1 μm, **c**:250 nm, **e**:25 nm, **f**:10 nm.

This micron-scale compartment, termed the “phage nucleus”, forms de novo upon infection and grows with the replicating viral DNA. Meanwhile, phage proteins are synthesized in the host cell cytoplasm. This includes PhuZ, a phage-encoded tubulin homolog that assembles into filaments that treadmill to transport empty capsid heads from their assembly sites at the host cell membrane to the surface of the phage nucleus. Capsids dock to the phage nucleus surface and are filled with viral DNA before detaching and completing assembly with phage tails. Mature particles are released by host cell lysis. In contrast to other characterized anti-restriction systems, the phage nucleus renders jumbo phages broadly immune to DNA-targeting host restriction systems, including CRISPR/Cas, throughout infection by serving as a physical barrier between the viral DNA and host nucleases (3, 4).

While we previously showed that the phage nuclear shell incorporates at least one abundant phage-encoded protein (2, 8), the overall composition and architecture of this structure are still largely unknown. Also unclear is how these phages address the challenges arising from the separation of transcription and translation, specifically the need for directional transport of mRNA out of the nucleus and transport of DNA processing enzymes into the phage nucleus (2). Finally, how genomic DNA produced in the nucleus is packaged into phage capsids assembled in the cytosol is also unknown (9).

## Results

### The in-cell molecular architecture of the phage nuclear shell

To gain insight into the native architecture of the phage nuclear shell, we performed focused ion-beam milling coupled with cryo-electron tomography (cryoFIB-ET) of *Pseudomonas chlororaphis* 200-B cells (henceforth, *P. chlororaphis*) infected with jumbo phage 201φ2-1 at 50-60 minutes post infection (mpi) (**Fig. 1, SI Fig. 1**). The observed phage nuclei were pleomorphic compartments devoid of ribosomes, bounded by a ∼6 nm thick proteinaceous shell (**Fig. 1b-d, SI Fig. 1**). Close inspection of the compartment perimeter revealed repeating doublets of globular densities with ∼11.5 nm spacing, suggesting that the shell consists of a single layer of proteins in a repeating array (**Fig. 1e, SI Fig. 1j**). Furthermore, we occasionally captured face-on views that exhibit a square lattice of densities with the same repeat spacing, reinforcing the idea of a repeating array of protomers (**SI Fig. 1k**).

Using subtomogram analysis to average over low-curvature regions from eight separate 201φ2-1 nuclei, we obtained a ∼24 Å resolution reconstruction of the phage nuclear shell (**Fig. 1f,g, SI Fig. 2 & Table 2**). The reconstruction reveals the shell as a quasi-p4, or square, lattice (quasi because of variable curvature). The repeating unit is an 11.5 × 11.5 nm square tetramer approximately 6 nm thick, with internal four-fold rotational symmetry. These units form a square lattice with a second 4-fold rotational symmetry axis at the corner of each square unit and 2-fold axes on each side (the p442 wallpaper group; **Fig. 1h, SI Fig. 2c,d**). The four individual protomer densities within each 11.5 × 11.5 nm unit measure ∼6 × 6 × 7.5 nm, dimensions consistent with a ∼70 kDa protein. Thus, the phage nuclear shell appears to be predominantly composed of a single protein component arranged in a square lattice.

### The protein chimallin is the major component of the nuclear shell

We previously showed that the abundant and early-expressed 201φ2-1 protein gp105 becomes integrated into the nuclear shell (2). Furthermore, 201φ2-1 gp105 has a molecular weight of 69.5 kDa, consistent with the size of an individual protomer density from our in situ cryo-ET map. Along with the high apparent compositional homogeneity of the shell as observed by cryo-ET, these data support 201φ2-1 gp105 as the principal component of the phage nuclear shell. Homologs of 201φ2-1 gp105 are encoded by a large set of jumbo phage that infect diverse bacteria including Pseudomonas, Vibrio, Salmonella, and Escherichia coli, but these proteins bear no detectable sequence homology to any other proteins. Because of its role in protecting the phage genome against host defenses, we term this protein chimallin (ChmA) after the *chimalli*, a shield carried by ancient Aztec warriors (10).

To understand the structure and assembly mechanisms of chimallin, we expressed and purified 201φ2-1 chimallin from E. coli. Size-exclusion chromatography and multi-angle light scattering (SEC-MALS) of purified chimallin indicated a mixture of oligomeric states including monomers, well-defined assemblies of approximately 1.2 MDa, and larger heterogeneous species ranging from 4 to 13 MDa (**Fig. 2a**). Cryo-ET of the largest species revealed pleomorphic, closed compartments with near-identical morphology to the phage nuclear shell we observe in situ (**Fig. 2b**). Analysis of the smaller, more defined assemblies by cryo-ET revealed a near-homogeneous population of cubic assemblies with a diameter of ∼22 nm, and a minor population of rectangular assemblies with dimensions ∼22 × 33.5 nm (**Fig. 2c**).

**Figure 2.**
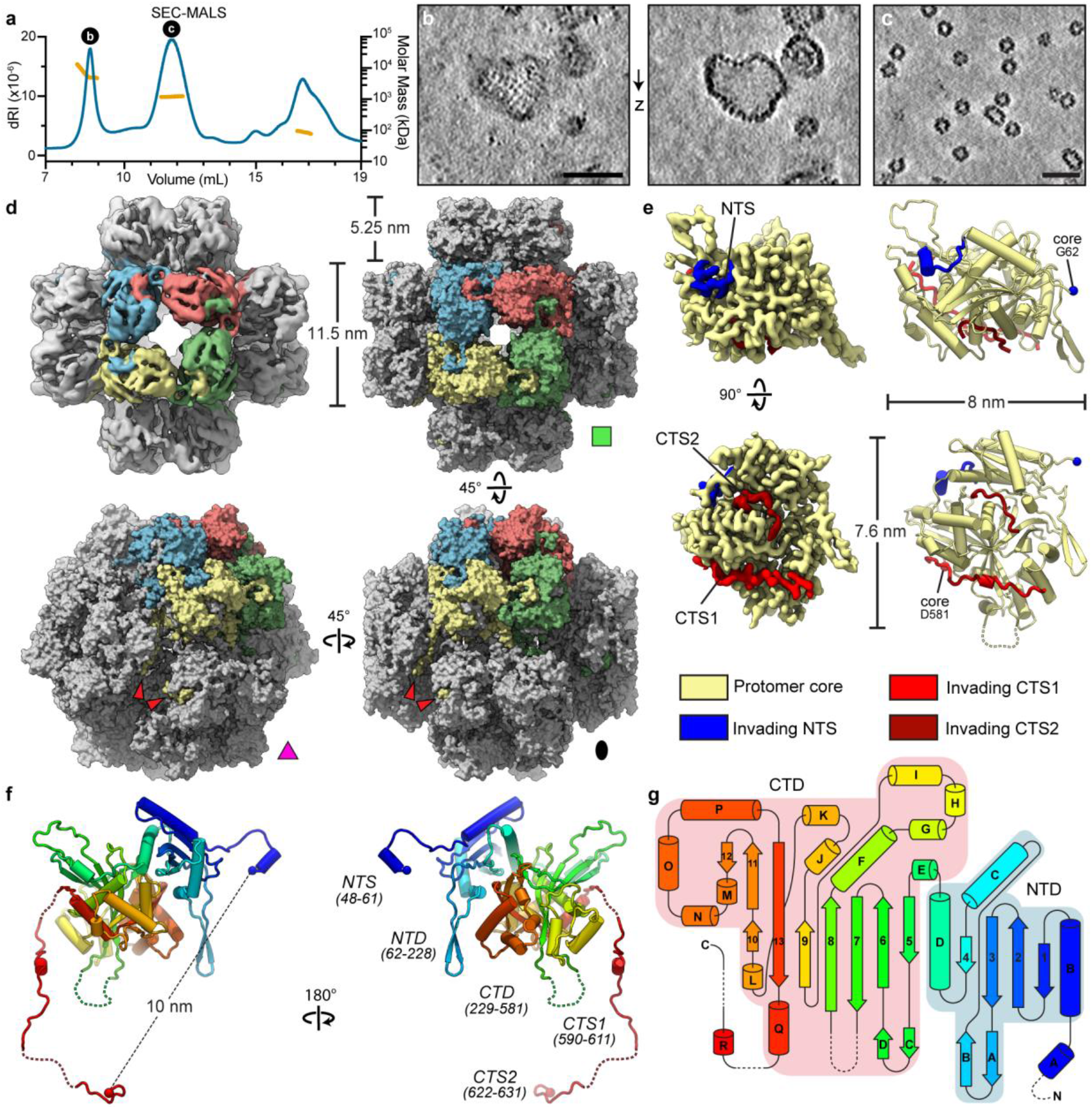
*In vitro* cryo-EM structure of the 201φ2-1 phage nuclear shell protein chimallin. **a**, Size-exclusion coupled to multi-angle light scattering (SEC-MALS) analysis of purified 201φ2-1 chimallin. From left to right, the measured molar mass of three peaks are 6.9 MDa (range from 4-13 MDa), 1.2 MDa, and 87 kDa. **b**,**c**, Z-slices from tomograms of samples from the correspondingly labelled SEC-MALS peaks in **a. d**, Top-left, O-symmetrized reconstruction of the chimallin cubic assembly viewed along the 4-fold axis. The protomers of one 4-fold face are colored. Top-right, surface representation of the chimallin cubic assembly model viewed along the 4-fold axis. Bottom-right and bottom-left, views of the model along the 2- and 3-fold axes, respectively. Red arrows point to the C-terminal segments of the yellow protomer. **e**, Localized asymmetric reconstruction of the chimallin protomer (left) and cartoon model (right). Invading N- and C-terminal segments from neighboring protomers are colored blue (NTS), red (CTS1), and burgundy (CTS2). Resolved core protomer termini are shown as spheres. **f**, Rainbow colored cartoon model of the Chimallin protomer conformation in the cubic assembly. Resolved N- and C-termini are shown as spheres. Domains and segments are labelled. Unresolved linkers are shown as dashed lines. **g**, A rainbow colored fold diagram of chimallin (blue at N-terminus, red at C-terminus) with ?-helices labelled alphabetically and *β*-strands labelled numerically. The N- and C-terminal domains are highlighted blue and red, respectively. Dashed lines indicate unresolved loops. Scale bars: **b**,**c**: 50 nm.

We next acquired cryo-electron microscopy (cryo-EM) data and performed single-particle analysis of the defined chimallin assemblies (**SI Figs. 3 & 4**). Two-dimensional class averages revealed that each cubic particle consists of six chimallin tetramers (24 protomers, 1.67 MDa) arranged to form a minimal closed compartment with apparent octahedral (O, 432) symmetry (**SI Fig. 3a**). Similarly, the minor population of ∼22 × 33.5 nm rectangular particles are assemblies of ten chimallin tetramers (40 protomers, 2.78 MDa) with apparent D4 symmetry (**SI Fig. 4**). Each tetrameric unit in these assemblies is an 11.5 × 11.5 nm square, in line with our in situ subtomogram analysis of the phage nuclear shell. Three-dimensional reconstruction of the cubic particles with enforced O symmetry resulted in a ∼4.4 Å density map with distorted features (**Fig. 2d**), likely arising from inherent plasticity of these assemblies breaking symmetry. Localized reconstruction of the square faces from each particle with C4 symmetry resulted in an improved density map at ∼3.4 Å. Further reduction of the structure to focus on an individual protomer resulted in the highest quality map at ∼3.1 Å, enabling atomic modelling of the chimallin protomer (**Fig. 2e,f, SI Tables 2-4**).

Chimallin folds into a compact two-domain core with extended N- and C-terminal segments (**Fig. 2f,g**). The N-terminal domain (residues 62-228) shows little structural homology to any characterized protein, adopting an α+*β* fold that is topologically similar only to an uncharacterized protein from *Enterococcus faecalis* (**SI Fig. 5a,b**). The C-terminal domain (residues 229-581) adopts a GCN5-related N-acetyltransferase fold most similar to that of *E. coli* AtaT and related tRNA acetylating toxins (PDB: 6AJM, C? RMSD 4.2 Å) (**SI Fig. 5c-e**) (11). While chimallin lacks the acetyltransferase active site residues of AtaT and related toxins, this structural similarity suggests that the jumbo phage nuclear shell may have evolved from a bacterial toxin-antitoxin system.

Atomic models of the C4-symmetric chimallin tetramer and quasi-O symmetric cubic assemblies reveal the molecular basis for chimallin self-assembly. The chimallin protomer map contains three interacting peptide segments from the N- and C-termini of neighboring protomers (**Fig. 2d-f**). Using the maps for the tetrameric face and full cubic assembly, we built models for the higher-order chimallin oligomers to understand the subunit interconnectivity. The N-terminal interacting segment (NTS, residues 48-61) of each protomer extends counterclockwise (as viewed from outside the cube) and docks against a neighboring protomer’s N-terminal domain within a given face of the cubic assembly, thus establishing intra-tetramer connections. Meanwhile, two extended segments of the C-terminus (CTS1: residues 590-611, and CTS2: residues 622-631) establish inter-tetramer interactions. While the linkers between the C-terminal domain and CTS1 (residues 582-589) and between CTS1 and CTS2 (residues 612-621) are unresolved in our maps, we could confidently infer the path of each protomer’s C-terminus within the cubic assembly. CTS1 extends from one protomer to a neighboring protomer positioned counter-clockwise around the three-fold symmetry axis of the cube, and CTS2 further extends counterclockwise to the third subunit around the same axis (**Fig. 2d**).

Notably, the binding of CTS1 to the chimallin C-terminal domain resembles the interaction of the antitoxin AtaR with the AtaT acetyltransferase toxin (**SI Fig. 5e**) (11), further hinting that the phage nuclear shell could have evolved from a bacterial toxin-antitoxin system. Compared to the tightly-packed chimallin tetramers mediated by the well-ordered NTS region, the length and flexibility of the linkers between the chimallin C-terminal domain, CTS1, and CTS2 suggest that flexible inter-tetramer packing enables chimallin to assemble into structures ranging from a flat sheet to the observed cubic assembly.

### Molecular basis for chimallin self-assembly & dynamics

To investigate the interconnectivity of chimallin protomers in the context of the phage nucleus, we docked copies of the high-resolution chimallin tetramer model into the in situ cryo-ET map. The chimallin tetramers fit well into our ∼24 Å resolution cryo-ET map of the shell without clashes and with an overall map-model correlation coefficient (CC) of 0.90 (**SI Fig. 6a**). To accommodate a flat sheet structure, the three-fold symmetry axis at each corner of the cubic assembly must be “unfolded” into a four-fold symmetry axis (**Fig. 3a**). Since CTS1 and CTS2 mediate interactions across this symmetry axis in the cubic assembly, this change requires that CTS1 rotate ∼55° relative to the chimallin protomer core, and that CTS2 rotate the same amount relative to CTS1 (**Fig. 3b**). In the resulting sheet, each chimallin C-terminus extends counterclockwise to contact the two neighboring subunits in the new four-fold symmetry axis at the corners of the tetrameric units. The distances spanned by each disordered linker are similar in the flat sheet and the cubic assembly (**Fig. 3b**). Thus, both N- and C-terminal interacting segments contribute to shell self-assembly, with the C-terminus in particular likely imparting significant structural plasticity to the phage shell while maintaining its overall integrity.

**Figure 3.**
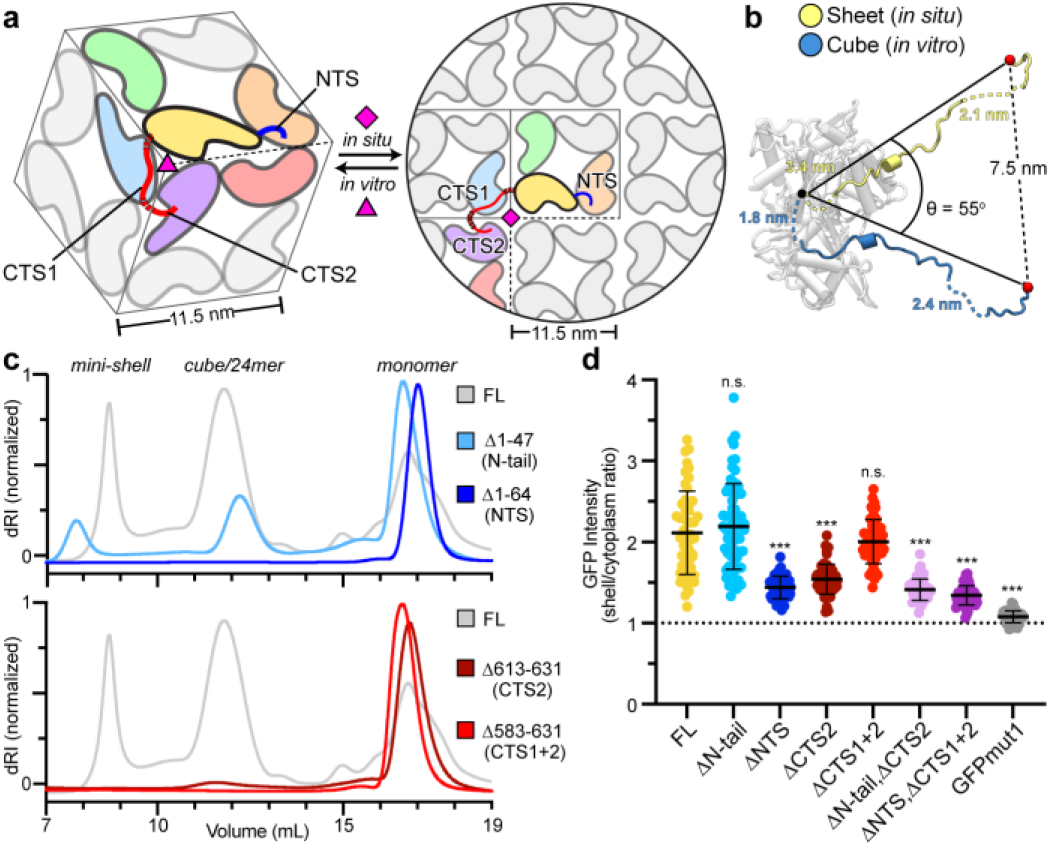
Flexibly attached N- and C-terminal segments mediate self-assembly of the chimallin shell. **a**, Relationship of 201φ2-1 chimallin protomer packing in the cubic/24mer assemblies (left) and flat sheet model (right). One protomer is colored yellow with its NTS in blue and CTS1/CTS2 in red. Protomers that interact directly with this central protomer are colored orange, green, blue, purple, and red. Non-interfacing protomers are shown in gray. Red arrows point to locations of unresolved linkers (red dashed lines), and pink symbols indicate 3- or 4-fold symmetry axes. **b**, Comparison of chimallin C-terminus conformation in the *in vitro* sheet (yellow) and *in situ* cube (blue) models. The distances spanned by each disordered segment (CTD-CTS1: residues 582-589; CTS1-CTS2: residues 612-621) in the two models are noted. **c**, SEC-MALS profiles of N- and C-terminal truncation mutants (ΔN-tail=Δ1-47; ΔNTS=Δ1-64; ΔCTS2=Δ613-631; ΔCTS1+2=Δ583-631). dRI: differential refractive index, See **SI Fig. 7** for molar mass measurements by SEC-MALS. **d**, Relative incorporation of GFPmut1-chimallin variants into the 201φ2-1 phage nuclear shell of infected *P. chlororaphis* cells. Incorporation is calculated as the ratio of GFP fluorescence/pixel in the shell vs. outside the shell (**SI Fig. 8 & Methods** for details). Data are the mean ± s.d. Statistical analysis performed was an unpaired t-test between a given variant and the full-length (FL, n = 67) protein ‘***’ indicates a calculated P-value < 0.0001: ΔN-tail (n = 51, P = 0.4131) ; ΔNTS (n = 53, P < 0.0001); ΔCTS1 (n = 54, P < 0.0001); ΔCTS1+2 (n = 50, P = 0.1884); ΔN-tail, ΔCTS2 (n = 58, P < 0.0001); ΔNTS, ΔCTS1+2 (n = 63, P < 0.0001); GFPmut1 (n = 60, P < 0.0001).

We next assessed the importance of the NTS and CTS regions for chimallin self-assembly both in vitro and in vivo. In vitro, deletion of either the NTS or CTS (CTS2 alone or CTS1+CTS2) completely disrupted self-assembly as measured by SEC-MALS (**Fig. 3c, SI Fig. 7**). We expressed the same truncations in phage 201φ2-1-infected P. chlororaphis cells, and measured incorporation of GFP-chimallin into the phage nuclear shell. Of note, targeted genetic knock-outs are currently not possible in nucleus-forming jumbo phage, thus these experiments were performed in the presence of abundant full-length chimallin produced by the jumbo phage, complicating the analysis. Nonetheless, we observed that deletion of the NTS, CTS, or both partially compromised incorporation of chimallin into the phage nuclear shell (**Fig. 3d, SI Fig. 8**). Overall, these data support the idea that the chimallin NTS and CTS regions are important for efficient shell assembly in infected cells. Moreover, the finding that even high-level expression of truncated protomers fails to disrupt the shell’s integrity underscores the extraordinary cooperativity of shell self-assembly.

### The nuclear shell balances flexibility & integrity

The phage nucleus shields the viral genome similar to the viral capsid. However, unlike viral capsids, chimallin does not tightly interact with the encapsulated DNA. Indeed, estimation of the electrostatics of chimallin indicate that both cytosolic and lumenal faces are negatively charged (**Fig. 4a,b**). This is in contrast to viral capsids which tend to have a positively-charged lumenal face to favorably interact with the negatively-charged DNA (12). The negative character of the phage nuclear shell likely mitigates interactions with the enclosed DNA, thereby keeping the genetic material accessible for transcription, replication, and capsid packaging.

**Figure 4.**
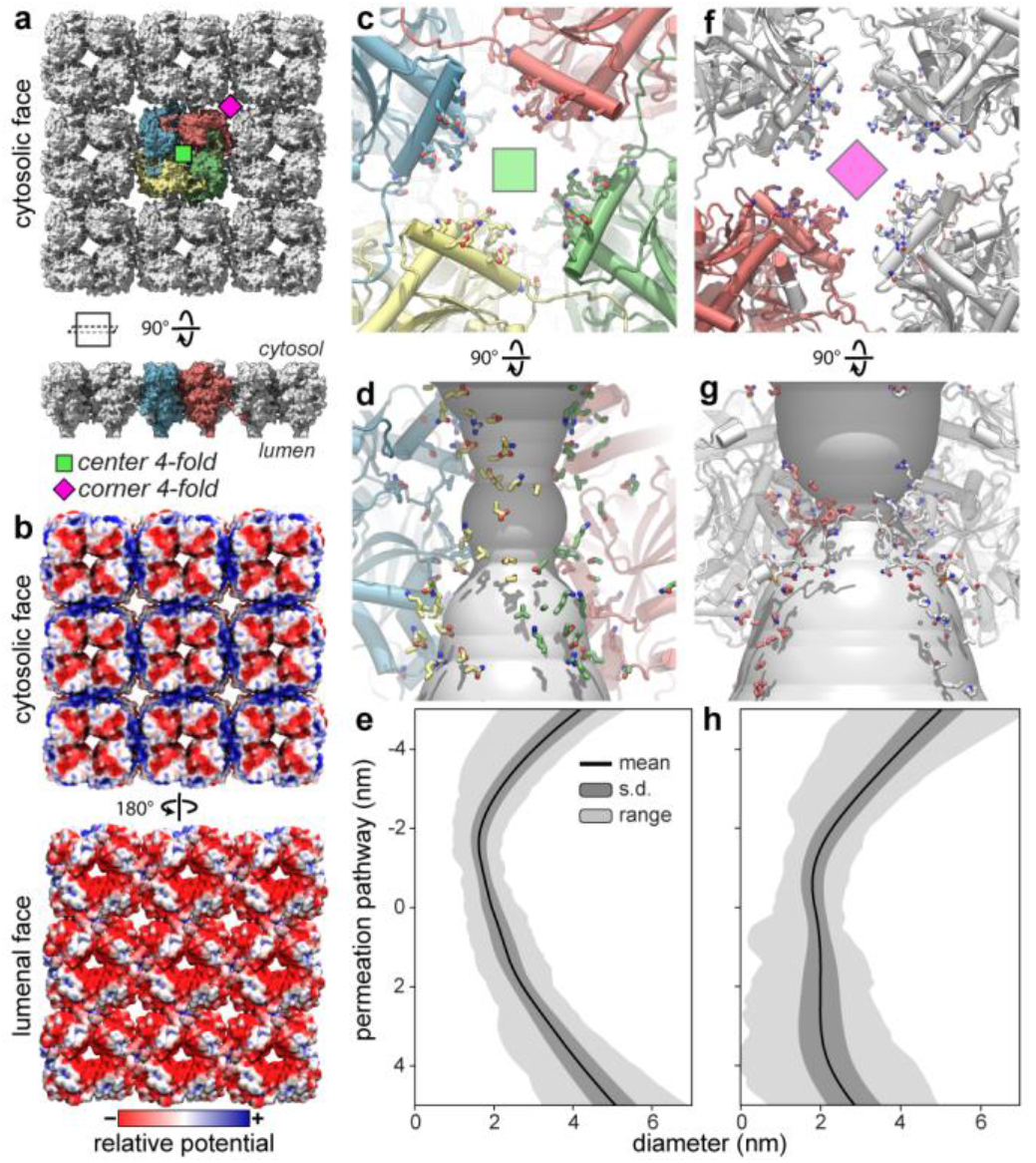
Properties of the chimallin lattice and four-fold pores. **a**, Surface model of the 3×3 chimallin tetramer lattice viewed from the cytosol and perpendicular slab view with the central tetramer colored. A ‘center’ four-fold is indicated by a green square and ‘corner’ four-fold by a magenta square. **b**, Surface model of the cytosolic and lumenal faces of the chimallin lattice colored by relative electrostatic potential. **c**,**d**, Cytosolic and side views respectively of the center four-fold pore cartoon model with pore-facing residues (**SI Table 8**) shown as sticks. The pore cavity for the chimallin lattice model prior to the molecular dynamics simulation shown as a gray surface. **e**, Representative (n = 9, **SI Fig. 10**) pore diameter profile for the center four-fold marked in a over the course of the 300-ns molecular dynamics simulation. The permeation pathway from top (negative values) to bottom (positive values) corresponds with cytosol to lumen. Solid black lines denote the mean diameter, dark gray shading +/- one standard deviation, and light grey shading the range. **f**,**g**, Same as **c**,**d** for the corner four-fold pore. h, Same as e for the corner four-fold marked in a (n = 4, **SI Fig. 11**)

Again, in contrast to viral capsids and other protein-based organelles such as bacterial microcompartments, which form regular assemblies with defined facets, the phage nuclear shell adopts a highly irregular morphology (**Fig. 1c, SI Fig. 1**). To identify the forms of conformational heterogeneity within the chimallin lattice, we performed an elastic network model analysis of a 3×3 tetramer chimallin sheet. This analysis not only indicated hinging along tetramer boundaries, which would lead to the cubic arrangement observed in vitro, but also hinging within a given tetramer (**SI Fig. 9, Videos 1 & 2**). Prompted by this analysis, we performed focused classification with our in situ data which revealed three distinct classes of the central chimallin tetramer (**SI Fig. 2e & 10**). The predominant class shows a flat sheet, while two minor classes show the central unit raised (convex) or lowered (concave) by ∼1 nm compared to surrounding units. Docking the tetramer model into maps representing concave, flat, and convex subpopulations indicated that the best fit is to the convex class (model-map CC = 0.93), and the worst fit is to the concave class (model-map CC = 0.88). Moreover, close inspection of the density representing a chimallin protomer in the different classes revealed that each protomer rotates ∼25° between the concave and convex classes, with the inner side of the tetramer pinching inward in the convex class (**SI Fig. 10d**). The tetramer model derived from the cubic assembly, which represents a highly convex state, shows a further ∼25° inward tilt of each protomer. These observations suggest that the chimallin tetramer itself is flexible, with the C-terminal domains on the shell’s inner face rotating inward in convex regions of the shell, and outward in concave regions. Thus, the morphology and flexibility of the phage nuclear shell likely derives from both intra- and inter-tetramer motions.

To analyse the inter-tetramer motions in the in situ data beyond the subtomograms that we extracted from relatively flat regions of the nuclear shell, we examined the flexibility between chimallin tetramers in *in situ* nuclear shells by calculating the curvature of an annotated surface in a tomogram and estimating the angle between tetramers. Our analysis showed that chimallin inter-tetramer conformations can range from slightly concave, i.e., -35° between neighboring tetramers, to highly convex, i.e., up to 80° between neighboring tetramers, close to the maximum bend of 90° observed in the purified sample (**SI Fig. 10e,f**). This enables the contorted shape observed in the cryo-ET data that presumably stems from addition of chimallin protomers to the lattice at multiple random locations, without distributing the strain throughout the lattice, rather than at a single privileged site.

### Pores within the chimallin lattice may accommodate transport of metabolites but not folded proteins

Like the eukaryotic nucleus, the phage nucleus separates transcription and translation (2). Thus, the phage nuclear shell must accommodate trafficking of mRNA out of the nucleus, and that of specific proteins into the nucleus. To determine whether the pores at the two four-fold symmetry axes of the chimallin lattice could serve as conduits for macromolecule transport, we used all-atom molecular dynamics to simulate the motions of a 3×3 flat sheet of chimallin tetramers (**Videos 3 & 4**). Assessing the variability in these pores through five separate 300 ns simulations, we found that the restrictive diameters of both the center and corner four-fold pores are ∼1.4 nm on average, varying throughout the simulations from ∼0 nm (i.e., closed) to as wide as 2.3 nm (**Fig. 4e,h, SI Fig. 11 & 12**).

These data strongly suggest that the pores are too small to accommodate most folded proteins. However, this pore size is sufficient to enable exchange of metabolites and potentially large enough to support export of single-stranded mRNA molecules. An intriguing model for mRNA export is suggested by prior findings on viral capsid-resident RNA polymerases, which physically dock onto the inner face of the capsid and extrude mRNA co-transcriptionally through ∼1.2-nm wide pores (13, 14). Notably, chimallin-encoding jumbo phages have been shown to encode distinctive multi-subunit RNA polymerases, suggesting co-evolution of chimallin and transcriptional machinery in this family (15). Further study will be required to determine the protein(s) and sites responsible for protein import, as well as whether mRNA is directly extruded through the chimallin lattice pores.

### Nuclear shell structure is conserved across jumbo phage

We recently discovered that the *E. coli* bacteriophage Goslar assembles a nuclear shell morphologically similar to those observed in the *Pseudomonas* phage 201φ2-1, φPA3, and φKZ (16). Goslar encodes a divergent homolog of 201φ2-1 chimallin (gp189), with 19.3% overall sequence identity between the two proteins (**Fig. 5a**). We performed cryoFIB-ET on *E. coli* cells infected with Goslar, followed by subtomogram analysis of phage nuclei (**SI Fig. 13 & 14**). The resulting ∼30 Å resolution reconstruction showed striking overall similarity to the structure of the 201φ2-1 nuclear shell, with a square grid of 11.5 × 11.5 nm units (**Fig. 5b, SI Fig. 14b,c**).

**Figure 5.**
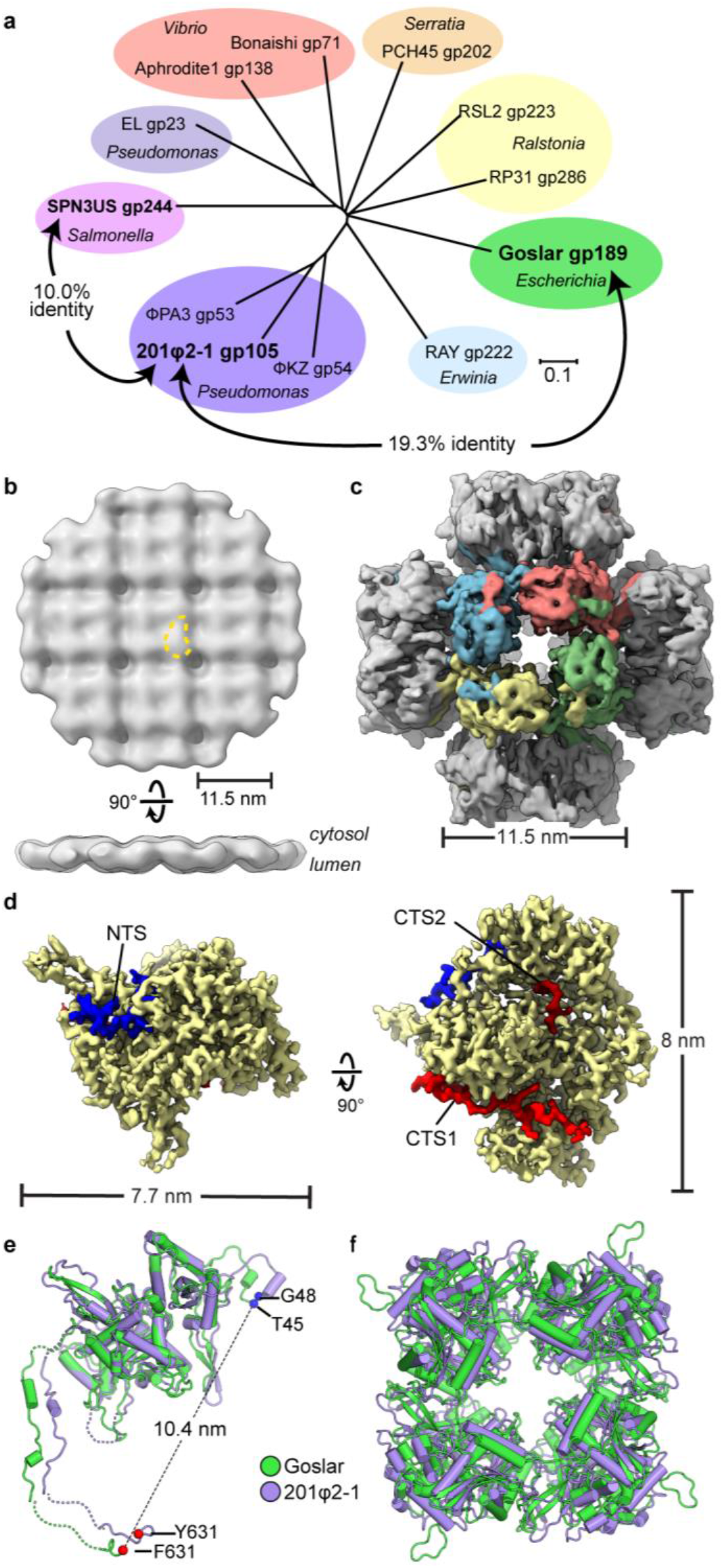
Structural conservation of chimallin in the distantly related *E. coli* jumbo phage Goslar. **a**, Unrooted phylogenetic tree of chimallin homologs. Homologs are listed as phage and gene product (gp) numbers (see **SI Table 9**). Groups based on proximity are colored and host genus labelled in italics (Scale bar = 0.1 substitutions/position). **b**, *In situ* subtomogram reconstruction of the Goslar chimallin shell. A comma-shaped protomer is marked by a yellow dashed outline and cytosolic and lumenal faces indicated. c, O-symmetrized map of the Goslar chimallin cubic assembly viewed along the 4-fold axis. **d**, Localized asymmetric reconstruction of the Goslar chimallin protomer (left) and cartoon model (right). Invading N- and C-terminal segments from neighboring protomers are colored blue (NTS), red (CTS1), and burgundy (CTS2). **e**,**f** Superposition of the Goslar (green) and 201φ2-1 (purple) coordinate models for the cube confirmation of the protomers **e** and the tetramers **f**. Resolved termini are shown as spheres for the protomers in **e**. The root mean square deviations are 1.8 Å and 4.8 Å for the protomers and tetramers, respectively.

We next purified Goslar chimallin and characterized its self-assembly by SEC-MALS. Like 201φ2-1 chimallin, Goslar chimallin forms a mixture of monomers, defined assemblies of ∼1.75 MDa, and large aggregates of ∼10 MDa (**SI Fig. 15a**). Cryo-EM analysis of the 1.75 MDa assemblies revealed cubic particles ∼22 nm in diameter, paralleling those formed by 201φ2-1 chimallin (**Fig. 5c, SI Fig. 15b**). We obtained a 4.2 Å resolution structure of the overall Goslar chimallin assembly, and used a similar localized reconstruction procedure as for the 201φ2-1 chimallin to obtain a 2.6 Å resolution structure of the Goslar chimallin tetramer, and a 2.3 Å resolution structure of a single protomer (**Fig. 5c,d, SI Fig. 12c-g**). Overall, Goslar chimallin shows high structural homology to 201φ2-1 chimallin despite the low overall sequence identity, with a C? RMSD of 1.8 Å within a protomer, and 4.8 Å over an entire C4-symmetric tetramer (**Fig. 5e,f**). The NTS, CTS1, and CTS2 segments of Goslar chimallin show near-identical interactions to neighboring protomers compared to 201φ2-1 chimallin with the exception of CTS2, which is shorter in Goslar than in 201φ2-1 and shows a distinct set of interactions (**SI Fig. 6e-g & Tables 5**,**6**).

Recently, negative-stain transmission electron microscopy of lysates from *Salmonella* cells infected with the jumbo phage SPN3US revealed an unidentified square lattice structure with a 13.5 nm periodicity (17). We identified a diverged chimallin homolog (gp244) in SPN3US that shares only 10% identity with 201φ2-1 chimallin (**Fig. 5a**). The similar dimensions and overall morphology of the SPN3US lattice compared to the 201φ2-1 and Goslar nuclear shells strongly suggest that these structures are composed of chimallin. Thus, together with our findings on 201φ2-1 and Goslar, these data show that despite extremely low sequence conservation, chimallin proteins from diverged jumbo phages exhibit high structural conservation at both the level of an individual protomer and overall nuclear shell architecture.

## Discussion

### Summary

Here, we detail the molecular architecture of the jumbo phage nuclear shell, a self-assembling, micron-scale proteinaceous compartment that segregates transcription and translation, largely excluding protein transport yet allowing selective protein import and mRNA export. We find that the shell is primarily composed of a single protein, termed chimallin, which self-assembles through its extended N- and C-termini into a closed compartment. The nuclear shell is a square quasi-lattice that effectively balances integrity and flexibility, erecting a physical barrier between the replicating phage genome and host defenses including restriction enzymes and CRISPR/Cas nucleases.

### Biogenesis of the phage nucleus & putative pre-nuclear shell compartment

Chimallin is the first and most abundant protein produced upon phage 201φ2-1 infection of a host cell (2). A key question is how chimallin specifically nucleates around the injected phage genome. Chimallin does not form a phage nucleus by itself when overexpressed in uninfected cells (2, 8), suggesting that the phage encodes nucleation factor(s) that promote assembly of the shell around its own genome. These factors may be produced alongside chimallin in the infected cell, or alternatively may be injected along with the phage DNA upon initial infection.

Thus, there is likely a window of time during the initial infection in which the phage genome is not protected by the nuclear shell. How does the phage protect itself during that time? We have consistently observed unidentified spherical bodies (USBs) in cells infected with either 201φ2-1 or Goslar, with internal density consistent with tightly packed DNA (**SI Fig. 16**). In 201φ2-1-infected cells, USBs average 201 nm in diameter (internal volume ∼4.25×10-3 μm3), while in Goslar-infected cells, they average 182 nm in diameter (internal volume ∼3.16×10-3 μm3). The internal volume of USBs in 201φ2-1-infected cells is 1.34x that of Goslar-infected cells, closely matching the 1.33x ratio between the genome sizes of two phages (317 kb for 201φ2-1 vs. 237 kb for Goslar). For both 201φ2-1 and Goslar, the internal volume of USBs is ∼3.3-4.4 times that of their capsids, suggesting that if each USB contains one phage genome, the DNA is less densely packed in USBs compared to the capsid. Using subtomogram analysis, we did not find USBs to have the same surface as the nuclear shell made of a single layer of chimallin; rather, the boundary’s density is more consistent with a lipid bilayer (**SI Fig. 16e-k**). Notably, similarly sized compartments have been observed by thin-section TEM of jumbo phage φKZ and SPN3US infecting *P. aeruginosa* and *Salmonella*, respectively (18, 19). Further work will be required to determine whether USBs represent unproductive infections that failed to pierce the inner membrane of the host, or instead represent a mechanism to protect a phage’s genome early in infection, prior to chimallin production and nuclear shell assembly.

### Phage nucleus growth and selective-permeability

Another unresolved question is how the phage nuclear shell grows as the phage genome replicates. We propose that shell growth is accomplished by incorporating soluble chimallin subunits into the lattice through a presumably isoenthalpic transfer of the N- and C-terminal segments that bind the chimallin interfaces on the lattice to the new chimallin subunits, effectively breaking and subsequently resealing the existing lattice. The high cooperativity of chimallin self-assembly, and the protein’s abundance in infected cells, likely ensures that host defense factors have little opportunity to access the replicating phage genome during its growth.

We and others have shown that specific phage proteins are actively imported into the phage nucleus, while other proteins – including host defense nucleases – are excluded (3, 4). Given that the pores of the chimallin lattice are not large enough to allow passage of most folded proteins, these data suggest that the phage encodes minor shell components that mediate specific, directional transport of proteins and potentially phage-encoded mRNAs through the protein barrier. Similarly, specific proteins either associated with or integrated into the shell likely mediate interactions with the phage-encoded tubulin homolog PhuZ to position the phage nucleus (2) and enable phage capsids to dock on the shell surface for genome packaging (9). Intriguingly, the prohead protease of the jumbo phage φKZ has been shown to cleave chimallin (gp54) between its C-terminal domain and CTS1 (**SI Fig. 17**) (20), suggesting that proteolytic processing of chimallin may contribute to capsid docking and filling by locally disrupting lattice integrity.

## Conclusion

The jumbo phage nucleus is a striking example of convergent evolution to solve a problem – isolation of a genome from the surrounding cell contents – previously thought to have evolved only once in the history of life. Here, we have shown how the phage-encoded chimallin protein (ChmA) cooperatively self-assembles into an effective nuclear-cytosolic barrier. This work sets the stage for future identification of minor shell components (ChmB, ChmC, etc.) that manage shell nucleation and growth, mediate nuclear-cytoplasmic transport, and direct key steps in the phage life cycle including cytoskeletal interactions and genome packaging. Finally, the structural elucidation of the principal component of the phage nuclear shell opens the possibility to the design of engineered protein-based compartments with sophisticated functions that span nanometer to micron scales.

## Acknowledgements

All electron microscopy data were collected at the UCSD Cryo-Electron Microscopy Facility, which was built and equipped with funds from UCSD and an initial gift from the Agouron Institute. We thank the UCSD Physics Computing for computational support. We thank V. Lam and J. Hutchings for advice on sample preparation and subtomogram averaging, respectively. We thank J. Krieger and I. Bahar at the University of Pittsburgh for discussions on elastic network models and expanding ProDy to accommodate the size of the chimallin sheet model. The authors thank members of the Pogliano, Villa, Corbett, and Amaro labs for helpful discussions and feedback. The authors acknowledge funding from the National Institutes of Health grants R01GM129245 (to J.P. and E.V.), R21AI148814 (to K.D.C.), and R01GM031749 (to J.A.M.), as well as from the National Science Foundation MRI grant NSF DBI 1920374 (to E.V.). T.L. is a Simons Foundation Awardee of the Life Sciences Research Foundation. C.S. is supported by a National Science Foundation Graduate Research Fellowship (DGE-1650112). E.V. is a Howard Hughes Medical Institute Investigator. Molecular graphics and analyses was performed in part with UCSF ChimeraX, developed by the Resource for Biocomputing, Visualization, and Informatics at the UCSF with support from National Institutes of Health R01-GM129325 and the Office of Cyber Infrastructure and Computational Biology, NIAID. This paper was typeset with a bioRxiv template by @Chrelli: www.github.com/chrelli/bi-oRxiv-word-template

## Author contributions

Conceptualization: TGL, AD, CS, REA, JP, KDC, EV; Methodology: TGL, AD, JP, KDC, EV; Validation: TGL, AD, CS, REA, JP, KDC, EV; Formal Analysis: TGL, AD, AMP, CS, REA, JP, KDC, EV; Investigation: TGL, AD, AMP, CS, YG, EE, SS, KK, EAB, EA; Data Curation: TGL, AD, CS; Writing - Original Draft: TGL, AD, CS, REA, KDC, EV; Writing- Review & Editing: AMP, YG, EE, SS, KK, EAB, EA, JAM, JP; Visualization: TGL, AD, AMP, CS, KDC, EV; Supervision: JAM, REA, JP, KDC, EV; Funding acquisition: TGL, CS, REA, JAM, JP, KDC, EV.

## Data availability

Cryo-EM density maps have been deposited in the Electron Microscopy Data Bank. Subtomogram averaging maps have the accession numbers: EMD-25221 (201φ2-1, consensus), EMD-25220 (201φ2-1, concave), EMD-25222 (201φ2-1, flat), EMD-25223 (201φ2-1, convex), EMD-25183 (P. chlororaphis, 70S), EMD-25229 (Goslar, consensus), EMD-25262 (Goslar, concave), EMD-25358 (Goslar, convex), EMD-25359 (APEC2248, 70S), EMD-25360 (APEC2248, 50S). Single-particle maps have the accession numbers: EMD-25393 (201φ2-1, O), EMD-25391 (201φ2-1, C4), EMD-25392 (201φ2-1, C1), EMD-25393 (201φ2-1, D4) EMD-25394 (Goslar, O), EMD-25395 (Goslar, C4), and EMD-25395 (Goslar, C1). Coordinate models have been deposited in the RCSB Protein Data Bank with the accession numbers 7SQQ (201φ2-1, O), 7SQR (201φ2-1, C4), 7SQS (201φ2-1 C1 monomer), 7SQT (Goslar, O), 7SQU (Goslar, C4), and 7SQV (Goslar, C1). Raw cryo-EM data have been deposited with the Electron Microscopy Public Image Archive with accession codes: EMPIAR-10859 (in situ 201φ2-1 tilt-series), EMPIAR-10860 (in situ Goslar tilt-series), EMPIAR-10862 (in vitro 201φ2-1 frame-series), and EMPIAR-10863 (in vitro Goslar frame-series). All other data are available upon request to the corresponding author(s).

## Materials and Methods

### Bacterial and phage growth conditions

*Pseudomonas chlororaphis* 200-B was grown on HA media at 30°C as previously described. 201φ2-1 lysates were collected as previously described with minor modifications (2). Briefly, 0.5 mL from a dense *P. chlororaphis* culture grown in HA liquid media (HA with no agar added) was infected with 10 μL serial dilutions of high titer 201φ2-1 lysate, incubated for 15 minutes at room temperature, mixed with 4.5 mL of HA 0.35% top agar, poured over HA plates, and incubated overnight at 30°C. Then, 5 mL phage buffer was added to web lysis plates the following day and incubated at room temperature for 5 hours. The phage lysate was collected by aspiration, cell debris was pelleted by centrifugation at 3220 × g for 10 minutes, and the resulting clarified phage lysate was stored at 4°C. *Escherichia coli* strain APEC 2248 was obtained from the DSMZ in Germany and grown in LB media at 37°C. Goslar lysate was sent by Johannes Whittmann at the DSMZ and stored at 4°C.

### Plasmid construction and expression

The pHERD-30T plasmid was used for expressing GFP-tagged full-length and truncated chimallin in *P. chlororaphis*. All constructs were designed with GFPmut1 fused to the N-terminus and synthesized by GenScript. The plasmids were electroporated into *P. chlororaphis*, and 25 μg/mL gentamicin sulfate was used for selection of colonies.

### Live cell fluorescence microscopy and image analysis

*P. chlororaphis* cells were inoculated on imaging pads in welled microscope slides. The imaging pads were composed of 1% agarose, 25% HA broth, 25 μg/mL gentamicin sulfate, 2 μg/mL FM4-64, 0.2 μg/mL DAPI, and 1% arabinose to induce expression. The slides were incubated at 30°C for 3 hours in a humid chamber, and 10 μL of undiluted high-titer 201φ2-1 lysate (1011 pfu/mL) was added to the pads to infect the cells 60 minutes before imaging. Samples were imaged using the DeltaVision Elite deconvolution microscope (Applied Precision, Issaquah, WA, USA). Images were deconvolved using the aggressive algorithm in the DeltaVision softWoRx program.

All image analysis was performed on images prior to deconvolution. Average protein incorporation into the 201φ2-1 phage nucleus structure was determined using FIJI by measuring the mean gray value of the ring of GFP intensity that denotes the phage nucleus and the mean gray value of cytoplasmic GFP outside of this ring. The ratio of mean gray values of this ring GFP to cytoplasmic GFP was calculated as the average incorporation. Representative cells were chosen from each dataset, and a 3D graph of normalized GFP intensity was generated in MATLAB 2019a.

### Cryo-electron microscopy of *in situ* samples and image acquisition

For grid preparation of 201φ2-1 infections, host bacterial cells were infected on agarose pads as previously described (2) for 50-60 minutes. For grid preparation of Goslar infections, 10 agarose pads (1% agarose, 25% LB) were prepared in welled slides and spotted with 10 µl of APEC 2248 cells at an OD600 of ∼0.35 then incubated at 37°C for 1.5 hours in a humidor. 10µl of Goslar lysate from the DSMZ was added to each pad then incubated for another 1.5 hours until being removed to room temperature. Infected cells were collected by the addition of 25 µl of 25% LB to each pad and gentle scraping with the bottom of an eppendorf tube followed by aspiration. A portion of the collection was aliquoted, the remainder was centrifuged at 6000 × g for 45 seconds, resuspended with 0.25x volume of the supernatant, and a portion of that was diluted 1:1 in supernatant. Plunging of samples began 20-30 minutes after removal from 37°C which significantly slows infection progression.

A volume of 4-7 µl of cells were deposited on R2/1 Cu 200 grids (Quantifoil) that had been glow-discharged for 1 min at 0.19 mbar and 20 mA in a PELCO easiGlow device shortly before use. Grids were mounted in a custom-built manual plunging device (Max Planck Institute of Biochemistry, Germany) and excess liquid blotted with filter paper (Whatman #1) from the backside of the grid for 5-7 seconds prior to freezing in a 50:50 ethane:propane mixture (Airgas) cooled by liquid nitrogen.

Grids were mounted into modified Autogrids (Thermo Fisher Scientific, TFS) compatible with cryo-focus ion beam milling. Samples were loaded into an Aquilos 2 cryo-focused ion beam/scanning electron microscope (TFS) and milled to generate lamellae approximately ∼150-250 nm thick as previously described (21).

Lamellae were imaged using a Titan Krios G3 transmission electron microscope (TFS) operated at 300 kV configured for fringe-free illumination and equipped with a K2 directed electron detector (Gatan) mounted post Quantum 968 LS imaging filter (Gatan). The microscope was operated in EFTEM mode with a slit-width of 20 eV and using a 70 µm objective aperture. Automated data acquisition was performed using SerialEM-v3.8b11 (22) and all images were collected using the K2 in counting mode.

For lamellae of 201φ2-1-infected *P. chlororaphis*, tilt-series were acquired at a 3.46 Å pixel size over a nominal range of +/-51° in 3° steps with a grouping 2 using a dose-symmetric scheme (23) with a per-tilt fluence of 1.8 e-·Å^−2^ and total of about 120 e-·Å^−2^ per tilt-series. Nine tilt-series were acquired with a realized defocus range of -4.5 to -6 µm along the tilt-axis. An additional two data sets of six tilt-series each were collected at a pixel size of 4.27 Å with nominal tilt ranges of +/-50° and +/-60° in 2° steps with a grouping 2 using a dose-symmetric scheme with a per-tilt fluence of 1.8-2.0 e-·Å^−2^ and total of about 100-110 e-·Å^−2^ per tilt-series.

For lamellae of Goslar-infected APEC 2248, tilt-series were acquired at a 4.27 Å pixel size over a nominal range of +/-56° in 2° steps with a grouping 2 using a dose-symmetric scheme with a per-tilt fluence of 2.6 e-·Å^−2^ and total of about 150 e-·Å^−2^ per tilt-series. Twenty-one tilt-series were acquired with a realized defocus range of -5 to -6 µm along the tilt-axis

### Image processing and subtomogram analysis of *in situ* cryo-electron microscopy data

All tilt-series pre-processing was performed using Warp-v1.09 unless otherwise specified (24). Tilt-movies were corrected for whole-frame motion and aligned via patch-tracking using Etomo (IMOD-v4.10.28) (25). Tomograms were reconstructed with the deconvolution filter for visualization and manual picking in 3dmod (IMOD-v4.10.28). All subsequently reported resolution estimates are based on the 0.143-cutoff criterion of the Fourier shell correlations between masked, independently refined half-maps using high-resolution noise-substitution to mitigate masking artifacts (26).

First, for the 201φ1-infected *P. chlororaphis* dataset collected at 3.46 Å per pixel, subtomogram averaging of the P. chlororaphis host cell ribosomes was performed in order to improve initial tilt-series alignments using the recently developed multi-particle framework, M (27) (**SI Fig. 2a & 18**). A set of 400 particles were manually picked across the tomograms, extracted at 20 Å per pixel, and aligned in RELION-v3.1.1 to generate an initial reference (28, 29). This data-derived reference was used for template-matching against 20 Å per pixel tomograms at a sampling rate of 15°. Hits were curated in Cube (https://github.com/dtegunov/cube), ultimately resulting in 17,169 particle picks. The initial particle set was extracted at 10 Å per pixel and subjected to Class3D, after which 11,148 particles were selected for further analysis. Refine3D of the curated particle set reached the binned Nyquist limit of 20 Å. The refined particles were imported into M-v1.09 at 3.46 Å per pixel. Three iterations of refinement were performed starting with image-warp and particle poses, then incorporating refinement of stage angles and volume-warp, and finally including individual tilt-movie alignment. This procedure resulted in a ribosome reconstruction at an estimated resolution of about 11 Å. Further refinement of the particles in RELION yielded a reconstruction at an estimated resolution of about 10 Å. Neither additional attempts at 3D-classification nor multi-particle refinement lead to an improved ribosome reconstruction.

New tomograms were reconstructed at 20 Å per pixel using the ribosome alignment metadata. The perimeters of the 201φ1 phage nuclei were coarsely traced in these updated tomograms using 3dmod (30). Traces were converted into surface models using custom MATLAB (MathWorks, v2019a) scripts and built-in Dynamo-v1.1.514 functions (31). Points were sampled every 4 nm along the surface models, oriented normal to the surface (i.e., positive-Z towards the cytosol), and extracted from normalized, CTF-corrected tomograms at 10 Å per pixel in a 480 Å side-length box using Dynamo (31). Initial orientations of the 66,887 extracted particles were curated using the Place Object plugin (32) for UCSF-Chimera-v1.15 (33) and incorrectly oriented particles were manually flipped.

To generate an initial reference, a subset of 17,622 particles from two tomograms were subjected to reference-free alignment in Dynamo-v1.1.514 for several iterations. For this procedure, no point-group symmetry was enforced, alignment was limited to 40 Å by an ad hoc lowpass filter each interaction, and the out-of-plane searches were restricted to prevent flipping of sidedness. The alignment converged to yield a reconstruction conforming to an apparent square (p4, 442) lattice. Analysis of particle positions and orientations using Place Objects (32) and “neighbor plots” (34) were consistent with reconstructed average and indicated a spacing between approximately 11.5 nm between congruent 4-fold axes.

The initial reference was subsequently used to align the entire dataset for a single iteration in Dynamo. For this step, alignment was limited to 40 Å, the out-of-plane searches were restricted to prevent flipping of sidedness, C4 symmetry enforced, and a box-wide by 240 Å cylindrical alignment mask applied. Inspection of particle positions and orientations using Place Objects (32) and neighbor plots (34) were again consistent with a square lattice-like arrangement. To deal with the initial over-sampling, particle duplicates were identified as those within 9 nm center-to-center distance of another particle and the one with the lower cross-correlation to the reconstruction removed, which resulted in 21,165 retained particles. In addition, a geometry-based cleaning step was performed to remove particles with less than three neighbors within 10 to 13 nm, which resulted in 8,454 retained particles.

The curated particle set was split into approximately equal half-sets on a per-tomogram basis, converted to the STAR file format using the dynamo2m-v0.2.2 package (35), and re-extracted into a 480 Å side-length box at 5 Å per pixel in Warp for use in RELION. A round of Refine3D was performed using a 40 Å lowpass filtered reference, C4 symmetry, local-searches starting at 3.7°, and a box-wide soft-shape mask. This resulted in a reconstruction at an estimated resolution of 24 Å for the 8,454 particle set.

Classification without alignment was performed using a 320 Å spherical mask and C4 symmetry, to promote convergence from the relatively low particle count. This differentiated three distinct classes corresponding to “concave” (2,033 particles), “flat” (4,475 particles), and “convex” (945 particles) states of the central tetramer, along with a noisy class (1,001 particles). The particles corresponding to the interpretable classes were re-extracted into a 320 Å box at 5 Å per pixel and subjected to 3D auto-refinement as described for the consensus reconstruction. The estimated resolutions for the lowered, intermediate, and raised classes were 20 Å, 18 Å, and 23 Å, respectively. Refinement in M of either the consensus particle set or the three aforementioned classes above did not yield notable improvements in the reconstructions. This may be attributed to prior refinement of the tilt-series alignment, the most resolution-limiting factor, using the host ribosomes (27).

The twelve tilt-series of 201φ1-infected *P. chlororaphis* dataset collected at 4.27 Å per pixel were pre-processed and host ribosomes averaged as described above. The ribosome reconstruction from above was low-pass filtered to 40 Å and used for template-matching in Warp-v1.09, and curated in Cube to yield 47,469 particle positions. Ribosomes were extracted at 10 Å per pixel and subjected to reference-free 3D classification in RELION-v3.1.1 which resulted in 15,782 subtomograms. Masked auto-refinement resulted in a reconstruction with an estimated resolution of 28 Å. Refinement of tilt-series parameters in M improved the resolution to about 20 Å (not shown).

For the 201φ1 nucleus in the 4.27 Å per pixel dataset, we were unable to completely resolve the lattice register using either an *ab initio* reference or the reconstruction from above as determined by neighbor plots. Thus, this dataset was solely included in the analysis of the unidentified spherical bodies and not further subtomogram averaging.

The tilt-series of Goslar-infected APEC 2248 were pre-processed and host ribosomes averaged similarly as described above (**SI Fig. 18**). An initial host ribosome reference was generated from 400 randomly selected particles and used for template-matching in Warp-v1.09, and curated in Cube to yield 98,981 particle positions. Ribosomes were extracted at 10 Å per pixel and subjected to reference-free 3D classification in RELION-v3.1.1 which distinguished 70S and 50S classes containing 46,056 and 3,710 particles, respectively. Particles were re-extracted at 6 Å per pixel and subjected to 3D auto-refinement to yield 20 Å and 14 Å for the 50S and 70S classes, respectively. Refinement of tilt-series parameters in M, followed by an additional round of 3D auto-refinement at 4.27 Å per pixel resulted in 12 Å and 8.54 Å (Nyquist-limit of the data) for the 50S and 70S, respectively.

For the Goslar nucleus, the nuclei perimeters were traced from 6 tomograms and used to extract over-sampled points normal to the surface, which resulted in 42,416 initial particles. A subset of 10,512 particles were used to generate an ab initio reference in C1 as described above. The initial Goslar reference presented a similar spacing (∼11.5 nm) and apparent C4 symmetry as the 201φ1 reconstruction, however the average converged on the opposite 4-fold axis. For ease of subsequent analysis, the center of the Goslar initial reference was shifted to match 201φ1. Alignment of the entire dataset was performed as described above and distance-based cleaning post-alignment resulted in 4,501 particles for further processing. Alignment and reconstruction in RELION enforcing C4 symmetry resulted in a reconstruction with an estimated resolution of 27 Å. Similar to the 201φ1, 3D classification of the consensus refinement without alignment separated the data into classes in which the central tetramer appeared “concave’ (2,802) or “convex” (1,699). Refinement of these classes resulted in reconstruction with estimated resolutions of 20 Å and 25 Å for the lowered and raised classes, respectively.

### Analysis of unidentified spherical bodies

Unidentified spherical bodies (USBs) were manually identified and their maximal apparent diameters measured from their line intensity profiles in 20 Å per pixel tomograms using FIJI (36). In order to assess whether the surfaces of these compartments possessed an underlying structure, we attempted subtomogram analysis of the compartment surfaces essentially as described in (35, 37). We were unable to obtain a reconstruction exhibiting a regular underlying structure as assessed both visually and by neighbor plots. However, despite their differing exterior membrane, the interior density of the USBs is visually consistent with nucleic acid like that of the interior of the phage nucleus.

### Segmentation and visualization of in situ tomography data

Segmentation of host cell membranes and the phage nucleus perimeter was performed on 20 Å per pixel tomograms by first coarsely segmenting using TomoSegMemTV (38) followed by manual patching with Amira-v6.7 (TFS). For the purposes of segmentation, phage capsids, tails, PhuZ, and RecA-like particles were subjected to a coarse subtomogram averaging procedure using particles sampled at 20 Å per pixel. For capsids, all particles were manually picked. Reference-generation and alignment of capsids was performed by enforcing icosahedral symmetry with Relion-v3.1.1 (28, 29) (despite the capsids possessing C5 symmetry) in order to promote convergence from the low number of particles. For the phage tails, the start and end points along the filament axis were defined manually and used to seed over-sampled filament models in Dynamo-v1.1.514 (31, 39). An initial reference for the tail was generated using Dynamo-v1.1.514 from two full-length tails with clear polarity. The resulting reference displayed apparent C6 symmetry, which was enforced for the alignment of all tails from a given tomogram using Dynamo-v1.1.514 and Relion-v3.1.1. Similar to the phage tails, the PhuZ and RecA-like filaments were picked and refined but without enforcing symmetry. We do not report resolution claims for these averages and solely use them for display purposes in segmentations. Duplicate particles were removed and final averages were placed back in the reference-frame of their respective tomograms using dynamo_table_place. For clarity, a random subset of 500 ribosomes were selected for display in the segmentation.

### Surface curvature estimates from segmentation

The segmentation of the phage nucleus, depicted in Figure 1d, sampled at 2 nm/pixels, was used to estimate the principal curvature of the shell using PyCurv (40). PyCurv was run with default parameters and a hit radius of 3 pixels. For visualization purposes, the principal curvature values (κ1, κ2) were converted from a radius of curvature (r, nm^-1^) to an angle (θ, degrees) using the following formula:

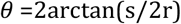

where r is the radius of curvature (inverse of κn) and s is for the side-length of the polygon circumscribed, taken as 5.75 nm.

### Protein expression and purification

Full-length Chimallin from bacteriophages 201φ2-1 (gp105; NCBI Accession YP_001956829.1) and Goslar (gp189; NCBI Accession YP_009820873.1) were cloned with an N-terminal TEV protease-cleavable His6-tag using UC Berkeley Macrolab vector 2-BT (Addgene #29666). Truncations and other modified constructs were cloned by PCR mutagenesis and isothermal assembly, and inserted into the same vector. Proteins were expressed in *E. coli* Rosetta2 pLysS (Novagen) by growing cells to A600=0.8, inducing expression with 0.3 mM IPTG, then growing cells at 20°C for 16-18 hours. Cells were harvested by centrifugation and resuspended in buffer A (50 mM Tris pH 7.5, 10 mM Imidazole, 300 mM NaCl, 10% glycerol, and 2 mM *β*-mercaptoethanol), then lysed by sonication and the lysate cleared by centrifugation. Protein was purified by Ni2+ affinity method. The purified proteins were centrifuged briefly to settle down the floating particles (visible large assemblies and aggregated proteins). The proteins were dialyzed into buffer B (20 mM Tris pH 7.5, 250 mM NaCl, 2 mM *β*-mercaptoethanol) and the N-terminal histidine tag was cleaved using TEV protease with overnight incubation at 4°C. The retrieved tagless proteins were further purified for homogeneity through Superose 6 Increase 10/300 GL column (Cytiva) in buffer B. The quality of purified proteins was verified by SDS-PAGE analysis.

For analysis by size exclusion chromatography coupled to multi-angle light scattering (SEC-MALS), a 100 μL sample of protein at 2 mg/mL was passed over a Superose 6 Increase 10/300 GL column (Cytiva) in buffer B. Light scattering and refractive index profiles were collected by miniDAWN TREOS and Optilab T-rEX detectors (Wyatt Technology), respectively, and molecular weight was calculated using ASTRA v. 8 software (Wyatt Technology).

Cryo-electron microscopy of in vitro samples and image acquisition For grid preparation, freshly purified recombinant 201φ2-1 chimallin was collected from size-exclusion chromatography (estimated concentration of 4 µM of the monomer, 0.3 mg/mL). Immediately prior to use, R2/2 Cu 300 grids (Quantifoil) were glow-discharged for 1 min at 0.19 mbar and 20 mA in a PELCO easiGlow device. Sample was applied to a grid as a 3.2 µL drop in the environmental chamber of a Vitrobot Mark IV (TFS) held at 16°C and 100% humidity. Upon application of the sample, the grid was blotted immediately with filter paper for 3 seconds prior to plunging into a 50:50 ethane:propane mixture cooled by liquid nitrogen. Grids were mounted into standard AutoGrids (TFS) for imaging. Grids for recombinant Goslar chimallin protein were prepared similarly, but with the modification that the sample was concentrated to approximately 33 µM of the monomer (2.5 mg/mL) prior to plunge-freezing. The void peaks from each purification were frozen similarly at the eluted concentration after dilution 1:1 with 6 nm BSA-tracer gold (Electron Microscopy Sciences).

All samples were imaged using a Titan Krios G3 transmission electron microscope (TFS) operated at 300 kV configured for fringe-free illumination and equipped with a K2 directed electron detector (Gatan) mounted post Quantum 968 LS imaging filter (Gatan). The microscope was operated in EFTEM mode with a slit-width of 20 eV and using a 70 µm objective aperture. Automated data acquisition was performed using SerialEM-v3.8b11(22) and all images were collected using the K2 in counting mode.

For the 201φ2-1 24-mer sample, tilt-series were acquired using a pixel size of 1.376 Å with a per-tilt fluence of 4.7 e-·Å^−2^ using a dose-symmetric scheme (23) from +/- 51° in 3° steps and a grouping 3, resulting in a fluence of 164.5 e-·Å^−2^ per tilt-series. In total 4 tilt-series were collected with a realized defocus of -2.5 to -4 µm along the tilt-axis. Movies for single-particle analysis were recorded at a pixel size of 1.075 Å with fluence of 42.6 e-·Å^−2^ distributed uniformly over 40 frames. Automated data acquisition was performed using image-shift with active beam-tilt compensation to acquire 9 movies per hole per stage movement. In total 4,192 movies were acquired with a realized defocus range of -0.1 to -1.5 µm.

For the Goslar 24mer sample, movies for single-particle analysis were recorded at a pixel size of 0.8452 Å with fluence of 40 e-·Å^−2^ distributed uniformly over 44 frames. Automated data acquisition was again performed using image-shift with active beam-tilt compensation to acquire 10 movies per hole per stage movement. In total, 3921 movies were acquired with a realized defocus range of -0.1 to -1.5 µm.

For void peak samples, tilt-series were acquired similarly to that of the 201φ2-1 24mer sample but using a pixel size of 1.752 Å and tilt-range of +/- 60°.

### Image processing of *in vitro* cryo-electron microscopy data

All movie pre-processing was performed using Warp-v1.09 unless otherwise specified (24). Tilt-movies of the 201φ2-1 chimallin were corrected for whole-frame motion and aligned via patch-tracking using Etomo (IMOD-v4.10.28) (25). Tomograms were reconstructed with the deconvolution filter for visualization and manual picking of subtomograms using 3dmod (IMOD-v4.10.28) (30). A total of 203 manually picked subtomograms and their corresponding 3D-CTF volumes were reconstructed with a 288 Å side-length. Subtomograms were aligned and averaged initially in C1 by reference-free refinement as implemented in RELION-v3.1.1 (28, 29) to an estimated resolution of 22 Å. The C1 reconstruction displayed features consistent with a cubic assembly of the 201φ2-1 chimallin protomers. Thus, an additional round of refinement using the C1 reconstruction as a reference and enforcing O point-group symmetry improved the estimated resolution to 18 Å.

For the single-particle 201φ2-1 chimallin data, movies were motion-corrected with exposure-weighting and initial CTF parameters estimated using 5×5 grids. Micrographs were culled by thresholding for an estimated defocus in the range of 0.3-1.5 µm and CTF-fit resolutions better than 6 Å resulting in 4,098 micrographs for further processing. An initial set of 140,782 particle positions were picked with BoxNet2 (Warp-v1.09) (24) using a model re-trained on 20 manually curated micrographs and using a threshold of 0.95. Particle images were extracted using a 396 Å side-length. All further processing was performed using RELION-v3.1.1 (29) unless otherwise specified. A single round of reference-free 2D-classification was performed and the 128,798 particle images assigned to the averages displaying internal features were selected for further processing. At this stage, analysis of the 2D averages suggested the presence of 4-, 3-, and 2-fold symmetry axes, consistent with a cubic arrangement of the chimallin protomers in the particles. Thus, we subjected the particle images to 3D-refinement using the subtomogram average obtained from above as an initial reference lowpass filtered to 35 Å and O point-group symmetry enforced, which resulted in a reconstruction at an estimated resolution of 4.2 Å.

However, the reconstruction did not display features consistent with this resolution estimate (e.g., *β*-strands were not separated). The high apparent point-group symmetry and distribution of 2D class averages did not support the inflated resolution being due to a preferred orientation. Partitioning particles into half-sets by micrograph did not change the estimated resolution of reconstruction, indicating the inflated estimate was not due to splitting identical or adjacent particles across the half-sets. In addition, extensive 3D-classification with and without symmetry enforced did not yield distinct classes. Therefore, the possibility of quasi-symmetry was investigated by performing localized reconstructions of sub-structures within the particles (13). To reduce computational burden, the apparent O symmetry was first partially expanded to C4 using relion_particle_symmetry_expand to fully expand to C1 before removing redundant image replicates (noting that redundant views of the 4-fold axes possess the same last two Euler angles) to yield 772,788 sub-particles. Refinement of the partially expanded particles while enforcing C4 point-group symmetry and using a soft shape mask resulted in a reconstruction with an estimated resolution of 3.6 Å with notably improved features. Re-centering and re-extraction using a 245 Å side-length followed by refinement improved the estimated resolution to 3.6 Å. CTF refinement (41) (per particle defocus, per micrograph astigmatism, beam-tilt, and trefoil) and Bayesian polishing(42) successively improved the resolution further to 3.5 Å and 3.4 Å, respectively. A round of 2D-classification without alignment was performed to remove particles assigned to empty or poorly resolved classes, which yielded a set of 664,363 sub-particles and no change in the estimated resolution upon re-running 3D-refinement.

Although the reconstruction substantially improved through this procedure, the C4 map still exhibited distorted density (e.g., elongated helices). Attempts at 3D-classification did not separate distinct classes. Thus, a localized reconstruction was performed focused on the individual chimallin protomer in C1. Again, to reduce computational burden, before expanding the symmetry to C1 another round of Bayesian polishing was performed in which the sub-particle images were extracted using a 354 Å side-length and premultiplied by their CTF before cropping in real-space to a 200 Å side-length. After another round of 3D-refinement enforcing C4 point-group symmetry, the data was expanded to C1 which resulted in 2,657,452 sub-particles and refined to an estimated resolution of 3.3 Å. The Bayesian polishing job was re-run to extract sub-particles at the full box size and without premultiplication by their CTF for import into cryoSPARC-v3.2 (43). A single round of local non-uniform refinement (44) was performed in C1 using a user-supplied static mask, marginalization, and FSC noise-substitution options, which lead to a final reconstruction of the 201φ2-1 chimallin monomer at an estimated resolution of 3.1 Å.

The Goslar chimallin single-particle data were pre-processed similarly to the 201φ2-1 chimallin data, which after initial thresholding resulted in 2889 micrographs for further processing. Initial particle positions were identified using the 201φ2-1 chimallin-trained BoxNet2 (Warp-v1.09) (24, 44) model with a threshold of 0.1, which resulted in 289,387 picks. Particles were extracted using a 400 Å side-length and subjected to iterative rounds of 2D-classification and sub-selection, which resulted in 78,532 particles used for initial 3D-refinement. The Goslar chimallin particles exhibited the same quasi-symmetry as the 201φ2-1 chimallin described above, thus were processed using the same localized reconstruction procedure. The quasi-O, quasi-C4, and C1 reconstructions yielded estimated resolutions of 4.0 Å, 2.6 Å, and 2.4 Å, respectively. The quasi-C4 and C1 reconstructions within RELION (29) were performed on particle images that were extracted using a 470 Å side-length, premultiplied by their CTF, and cropped in real-space to a 200 Å side-length. The final C1 reconstruction was performed in cryoSPARC-v3.2 (43) as described above, which led to a final reconstruction of the Goslar chimallin monomer at an estimated resolution of 2.3 Å from 1,407,340 sub-particle images.

All resolution estimates are based on the 0.143-cutoff criterion of the Fourier shell correlations between masked independently refined half-maps using high-resolution noise-substitution to mitigate masking artifacts (26). Local resolution estimates were computed using RELION with default parameters. Resolution anisotropy for the C1 reconstructions were assessed using the 3DFSC (45) web server which reported sphericity values of 0.963 and 0.994 for the 201φ2-1 and Goslar maps, respectively.

Void peak tilt-series were processed similarly to the 201φ2-1 24-mer tilt-series, but using the gold-fiducials for alignment instead of patch-tracking in Etomo (25).

### Coordinate model building and refinement

Initial monomer models were generated via the DeepTracer web server (46) followed by manual building in COOT-v0.9.1 (47) and subjected to real-space refinement in PHENIX-v1.19.2 (48). To generate tetramer models, monomer models were rigid-body docked into the C4 maps using UCSF Chimera-v1.15 (33) and the N-terminal segments joined to the appropriate protomer cores. To generate 24mer models, tetramer models were rigid-body docked into the hexahedral maps and the C-terminal segments reassigned to the appropriate protomer cores. To ensure robust refinement, tetramer and 24mer structures were refined with C4 or O non-crystallographic symmetry (NCS) constraints and reference-model restraints based on high-resolution monomer structures. Isotropic atomic displacement parameters were refined against the respective unsharpened maps. All models were validated using MolProbity (49) and EMRinger (50) (SI Table 2). EMRinger scores for the 201φ2-1 24mer, tetramer, and monomer models were 0.46, 2.86, and 2.39, respectively. EMRinger scores for the Goslar 24mer, tetramer, and monomer models were 0.92, 3.10, and 3.59, respectively.

### Interface analysis

Interface analysis for the cubic assemblies to identify interacting residues and to calculate buried surface area was performed using the ePISA-v1.52 (51) and CaPTURE (52) web servers.

### Nine-tetramer sheet modelling

Nine chimallin tetramers were arranged in a flat sheet (3×3) structure by fitting in the consensus subtomogram average. Assuming the unfolding of the cubical assembly to create a flat sheet structure, the interacting C-terminal segments in the corner three-fold axis were re-assigned in a four-fold symmetry axis to the corresponding protomer. The missing residues between the C-terminal domain and C-terminal segment were built manually in COOT-v0.9.1 ensuring no clash with other modeled atoms (taking the flat sheet model in consideration) (47). The missing loop region in a protomer (residues 307-319) was built using the DaReUS-Loop web server (53). This modeled chain (residues 45-612) was used to re-create the flat-sheet structure by applying symmetry. Finally in this flat sheet model, the protruding C-terminal segments of peripheral protomers were trimmed and twelve interacting segments in the periphery were included in the final model (48 chains).

### Protonation state assignment and electrostatics estimates

The electrostatic surface representation was generated with the APBS-v3.0.0 (54) using the AMBER99 force field (55) and a pH of 7.5 for assigning protonation states using PROPKA-v3.4.0 (56) through PDB2PQR-v3.4.0 (56, 57).

### Elastic Network Models

Elastic network models (58–60) are a subset of normal mode analysis (61, 62). Here we used anisotropic network models (ANM) (63) and Gaussian network models (GNM) (64–66). Both of these models simplify the protein structure into a series of nodes, with an internode potential energy function governing node motion. To look at it another way, each mode is an eigenvector whose corresponding eigenvalue is the frequency of that motion in the model; lower frequencies correspond to dynamics that best describe the structure’s intrinsic motions. ProDy (version 1.0) is a software program enabling calculation of ANM and GNM modes (67, 68), which we used in this study. We created 20,412 nodes for the ANM and GNM calculations, which is the largest number of nodes ever used in ProDy.

The five lowest frequency GNM modes accounted for 76% of the overall variance. Considering we do not need to use all ENM modes to capture the system’s dynamics (69) we selected these five GNM modes and the five lowest frequency ANM modes to use in our models. The GNM’s Kirchoff matrix was built with a pairwise interaction cutoff distance of 10 Å and a spring constant of 1.0, while the ANM’s Hessian matrix used a pairwise interaction cutoff distance of 15 Å and a spring constant of 1.0. The ANM structural ensemble movies used an RMSD difference of 25 Å from the original conformation to display the protein sheet’s flexibility.

### Molecular dynamics simulations

Simulations were performed using the 9-tetramer chimallin sheet model. This structure was protonated and placed in a waterbox through Amber’s tleap module (70). The system was neutralized with Na+ using a 12-6 ion model (71, 72). The CUDA version 10.1 implementation (73–75) of Amber 20 was used (70). The water model used was OPC (76) with the Amber 19ffsb force field (77). The resulting system, including the protein and waterbox, contained 1,729,704 atoms. Energy minimization was performed for a total of 10,000 cycles using a combination of steepest descent and conjugate gradient methods (73) while the heavy atoms were restrained with a force constant of 10.0 kcal/(mol × Å2). Next, the system was slowly heated from 10.0 K to 300.0 K over 4 ns before stabilizing at 300.0 K for the next 6 ns using the NVT ensemble with a Langevin thermostat with a friction coefficient (collision frequency) of γ=5.0 ps-1 and the heavy atoms restrained with a force constant of 1.0 kcal/(mol × Å 2). Equilibration was performed in the NPT ensemble for 20 ns, using a timestep of 2 fs and the SHAKE algorithm, constraining bonds involving hydrogens (78). The equilibration temperature was set at 300.0 K with a Langevin thermostat with a friction coefficient (collision frequency) of γ=1.0 ps-1 (79, 80) and the pressure set to 1 bar with a Berendsen barostat (81) with relaxation time constant *τ*=1.0 ps-1 and a heavy atom restraint with a force constant of 0.1 kcal/(mol × Å2). Periodic boundary conditions were enforced with the van der Waals interaction cutoff at 8 Å, while long-range interactions were treated with the Particle mesh Ewald algorithm (82). After equilibration, the system was cloned into five replicates for the production runs, still set at 300.0 K in the NPT ensemble. Each was run for 300 ns, resulting in 1.5 µs of total sampling.

The resulting molecular dynamics trajectories were analyzed through CPPTRAJ-v.25.6 (83) and MDTraj-v1.9.4(84).

### Pore Analysis

Pore annotation was performed using HOLE-v2.2005 (85) for visualization of the static dimensions and CHAP-v0.9.1 (86) was used for all other analyses. The free energy and solvent density plots were averaged between physiologically identical pores across all simulation replicates. The intertetramer (“corner four-fold”) pore in the upper-left quadrant contained two frames that caused CHAP to crash; these frames were removed before averaging after consultation with the CHAP developers. Considering we still averaged 1502 frames × 4 pores - 2 bad frames = 6006 frames for the intertetramer pores, we do not feel that this removal causes any difference in our conclusions.

### Structure visualization and figure generation

Density maps, coordinate models, and simulation trajectories were visualized and figures generated with PyMOL-v2.5 (Schrödinger L., DeLano W., 2021), UCSF Chimera-v1.15 (33), ChimeraX-v1.2.5 (87), and VMD-1.9.4a35 (88).

---

**SI Figure 1.**
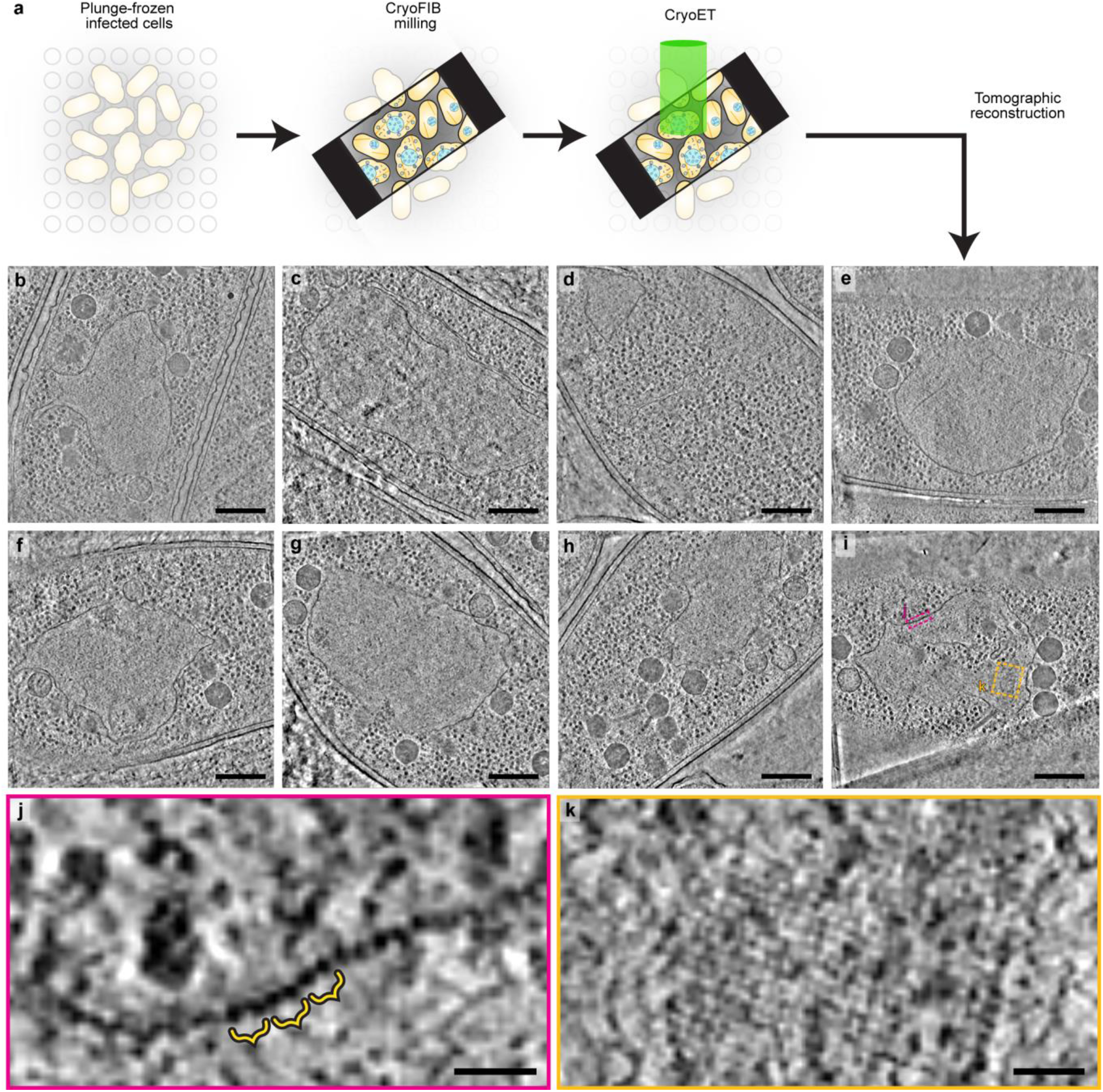
*In situ* cryoFIB-ET of 201φ2-1-infected *P. chlororaphis* cells. **a**, Schematic of cryoFIB-ET workflow. **b-i**, Slices of the eight 201φ2-1 nucleus-containing tomograms used in this study. **j**, Enlarged view of the corresponding colored boxed region in **i**. Exemplar doublets indicated by yellow brace. **k**, Enlarged view of the correspondingly colored boxed region in **i** which shows a square mesh-like texture corresponding to the square lattice. Scale bars: **b-i**: 250 nm, **j**,**k**: 25 nm.

**SI Figure 2.**
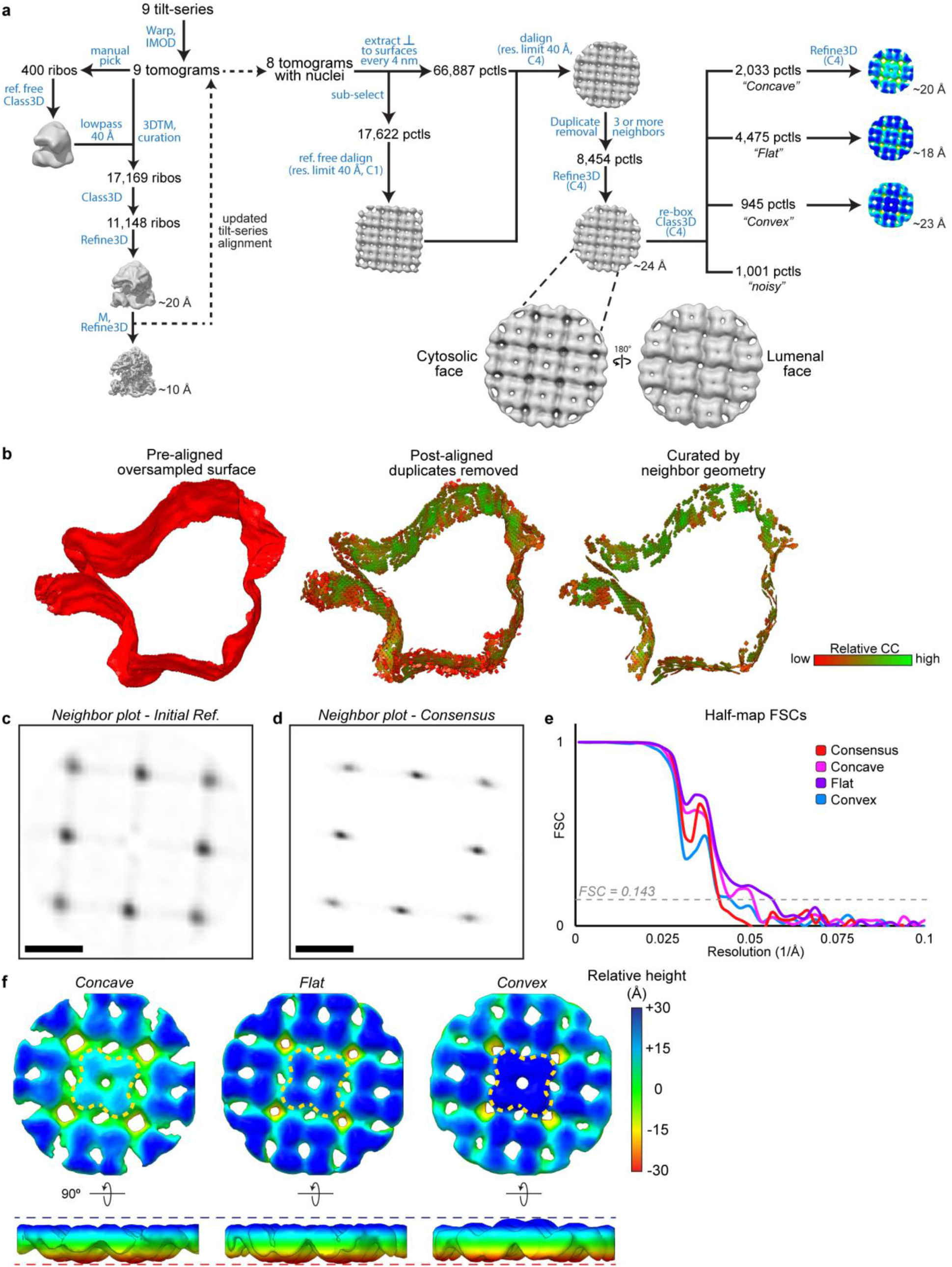
*In situ* subtomogram averaging of 201φ2-1 chimallinA in the context of the phage nucleus. **a**, Schematic of the subtomogram averaging workflow. Enlarged views of the consensus average with cytosolic and lumeal faces indicated. **b**, Example over-sampling and subtomogram curation strategy using lattice plots for the tomogram shown in Figure 1c. **c**, Neighbor plot of the initial, asymmetrically aligned reference. Inset scale bar is 10 nm. **d**, Neighbor plot of the symmetrized consensus refinement. **e**, Half-map FSC curves for the 201φ2-1 subtomogram reconstructions. **f**, Enlarged views of the resolved classes colored by relative height. The central tetramer is denoted by a yellow, dashed line for each class.

**SI Figure 3.**
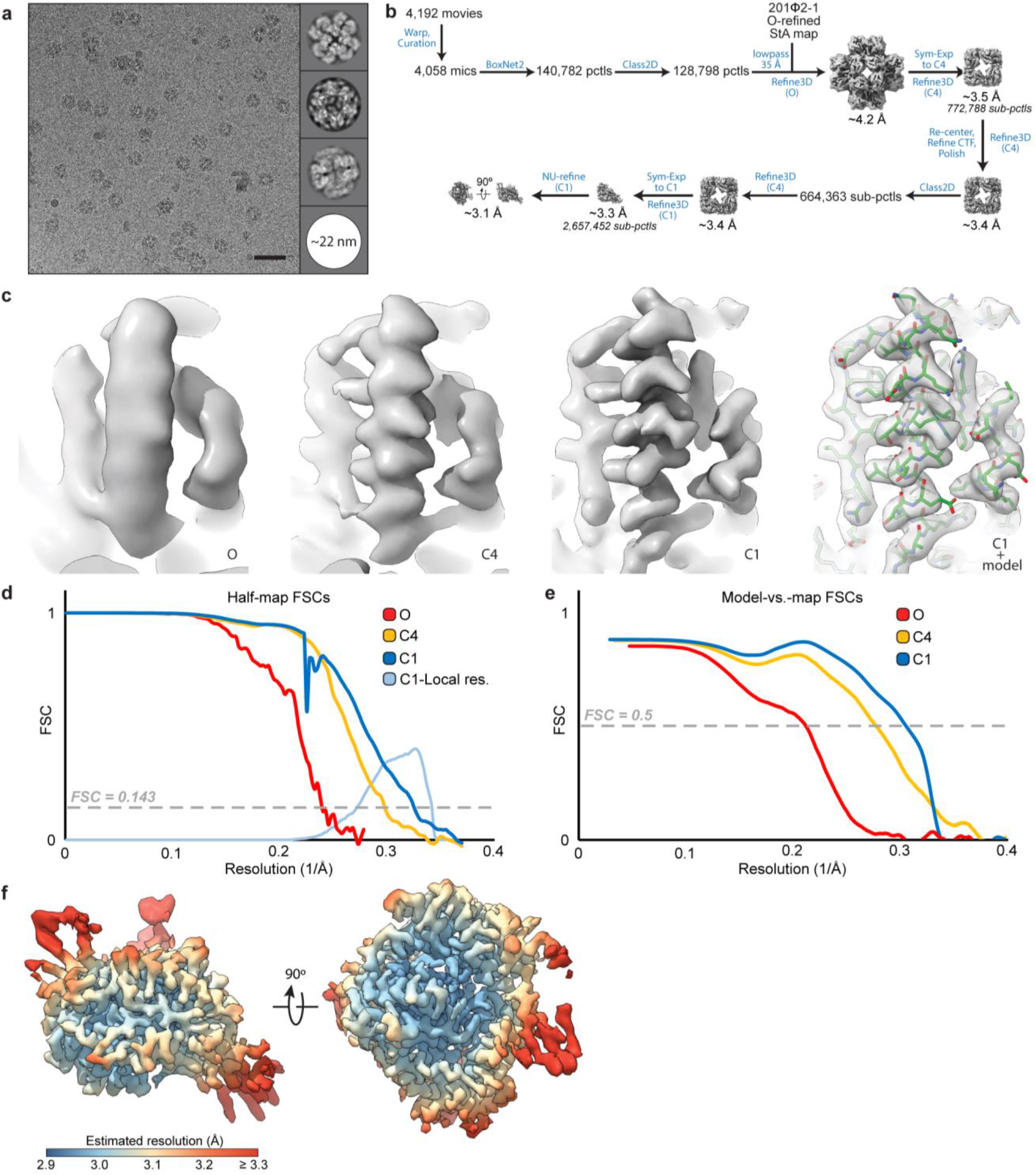
Single-particle reconstruction of the in vitro 201φ2-1 chimallinA cubic assembly. **a**, Exemplar micrograph and 2D class averages. Scale bar is 50 nm. **b**, Schematic of the localized reconstruction workflow. **c**, Unsharpened density map views centered on helix B (residues 68-84) at progressive stages of the localized reconstruction process. Final view of the C1 map shown with a fitted coordinate model. **d**, FSC curves for the half-maps at progressive stages of the localized reconstruction process (red, yellow, and blue), histogram of local resolution estimates for the C1 reconstruction (light blue). **f**, C1 reconstruction filtered and colored by local resolution.

**SI Figure 4.**
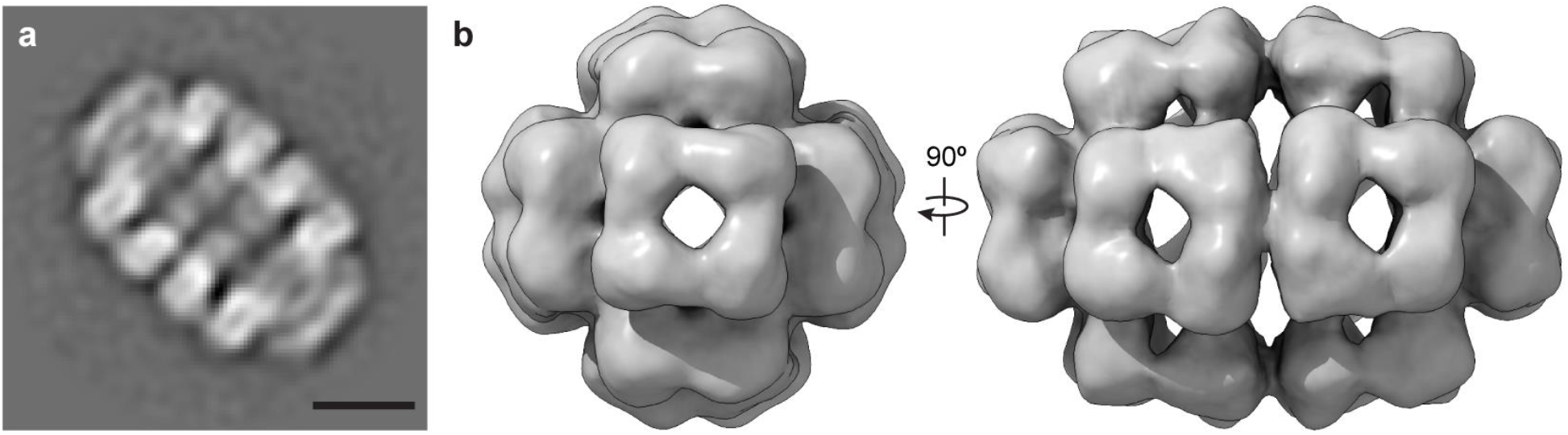
Single-particle reconstruction of the 201φ2-1 chimallinA elongated, quasi-D4 *in vitro* assembly. **a**, 2D class average of the minor (517 particles) species of elongated, quasi-D4 assemblies. Scale bar is 10 nm. **b**, Orthogonal views of the D4-symmeterized single particle reconstruction.

**SI Figure 5.**
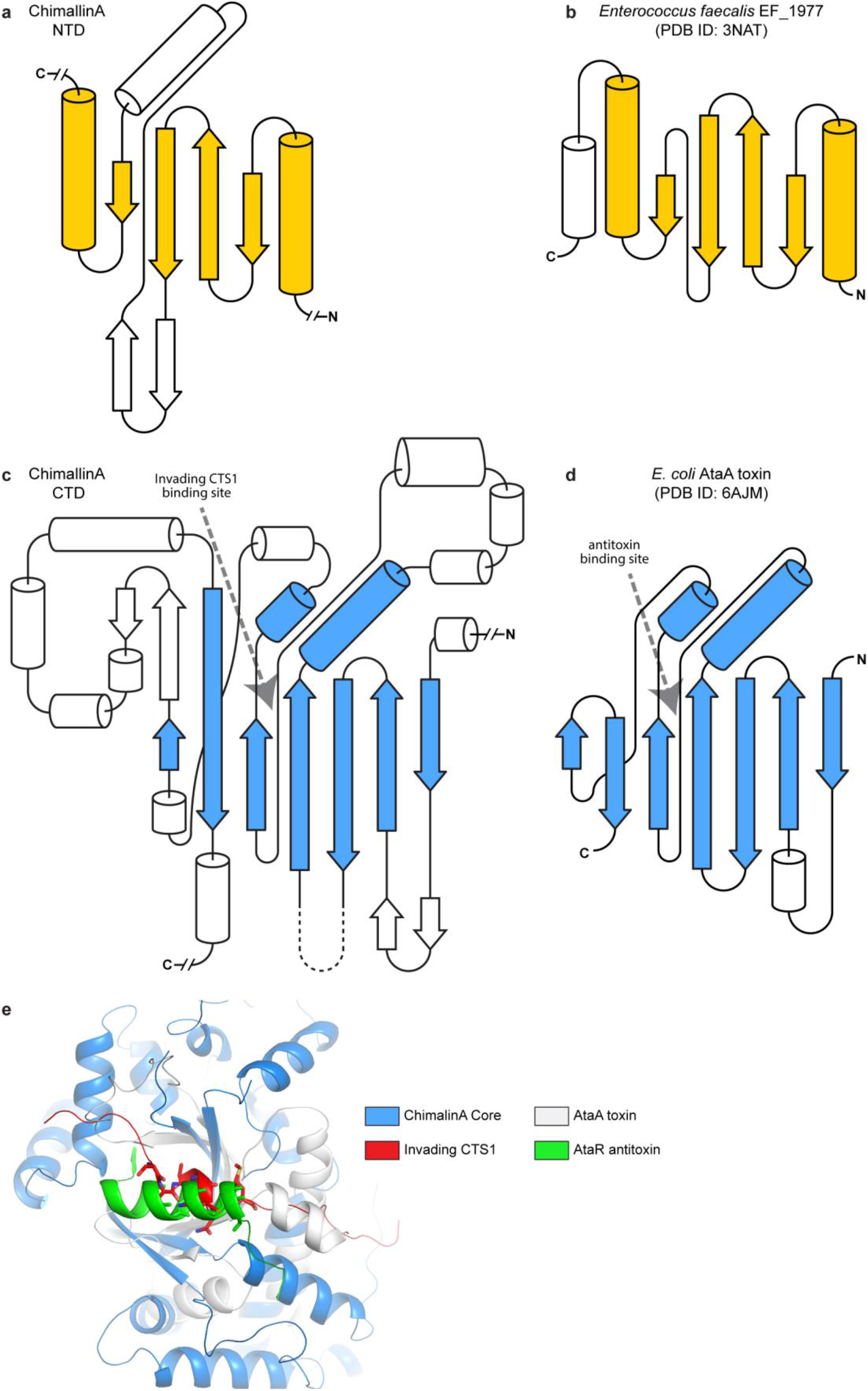
Partial homology of chimallinA fold topology to known structures. **a**, Topology of the 201φ2-1 chimallinA N-terminal domain (NTD, residues 62-228). **b**, Topology of E. faecalis EF_1977 (PDB ID: 3NAT), the closest structural relative of the chimallinA NTD. The root mean square deviation (RMSD) between chimallinA NTD and 3NAT coordinate models is 4.6 Å. Homologous secondary structure elements are colored in yellow. **c**, Topology of the 201φ2-1 chimallin C-terminal domain (CTD, residues 229-581). **d**, Topology of the E. coli AtaT tRNA-acetylating toxin (PDB ID: 6AJM) (11). The root mean square deviation (RMSD) between chimallinA CTD and 6AJM coordinate models is 4.2 Å. Homologous secondary structure elements are colored in blue. e, Structural overlay of the chimallinA CTD (blue) and AtaT (white; PDB ID 6AJM), showing the similarity in binding site for the chimallinA CTS1 segment (red) and the antitoxin AtaR (green).

**SI Figure 6.**
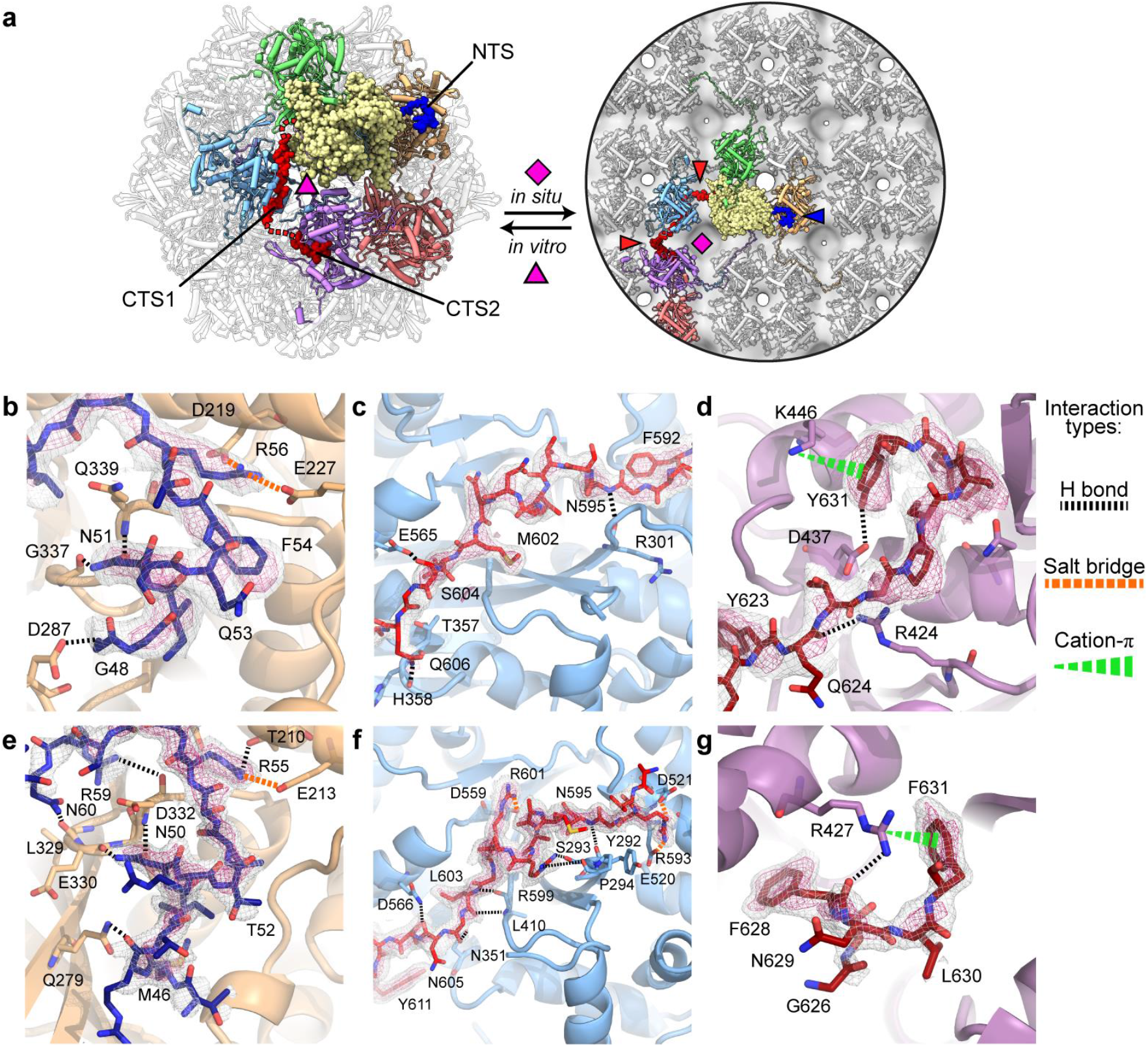
Interactions of NTS and CTS segments with the chimallinA core. **a**, Relationship of 201φ2-1 chimallin protomer packing in the cube (left) and flat sheet model (right). One protomer is shown as spheres and colored yellow with its NTS in blue and CTS1/CTS2 in red. Protomers that interact directly with this central protomer are colored. Non-interfacing protomers are in white. The flat sheet model is docked within the 201φ2-1 consensus subtomogram average map shown as transparent grey. Red arrows point to locations of unresolved linkers (red dashed lines), and pink symbols indicate 3- or 4-fold symmetry axes. **b-d**, Close-ups of the 201φ2- 1 coordinate model around the binding sites for NTS (**b**), CTS1 (**c**), and CTS2 (**d**). (**e-g**) Close-ups of the Goslar coordinate model around the binding sites for NTS (**e**), CTS1 (**f**), and CTS2 (**g**). For all panels, cryo-EM density map is shown as a mesh at high (pink) and low (grey) contours. Polar interactions are depicted by the symbols indicated in the key at the far right.

**SI Figure 7.**
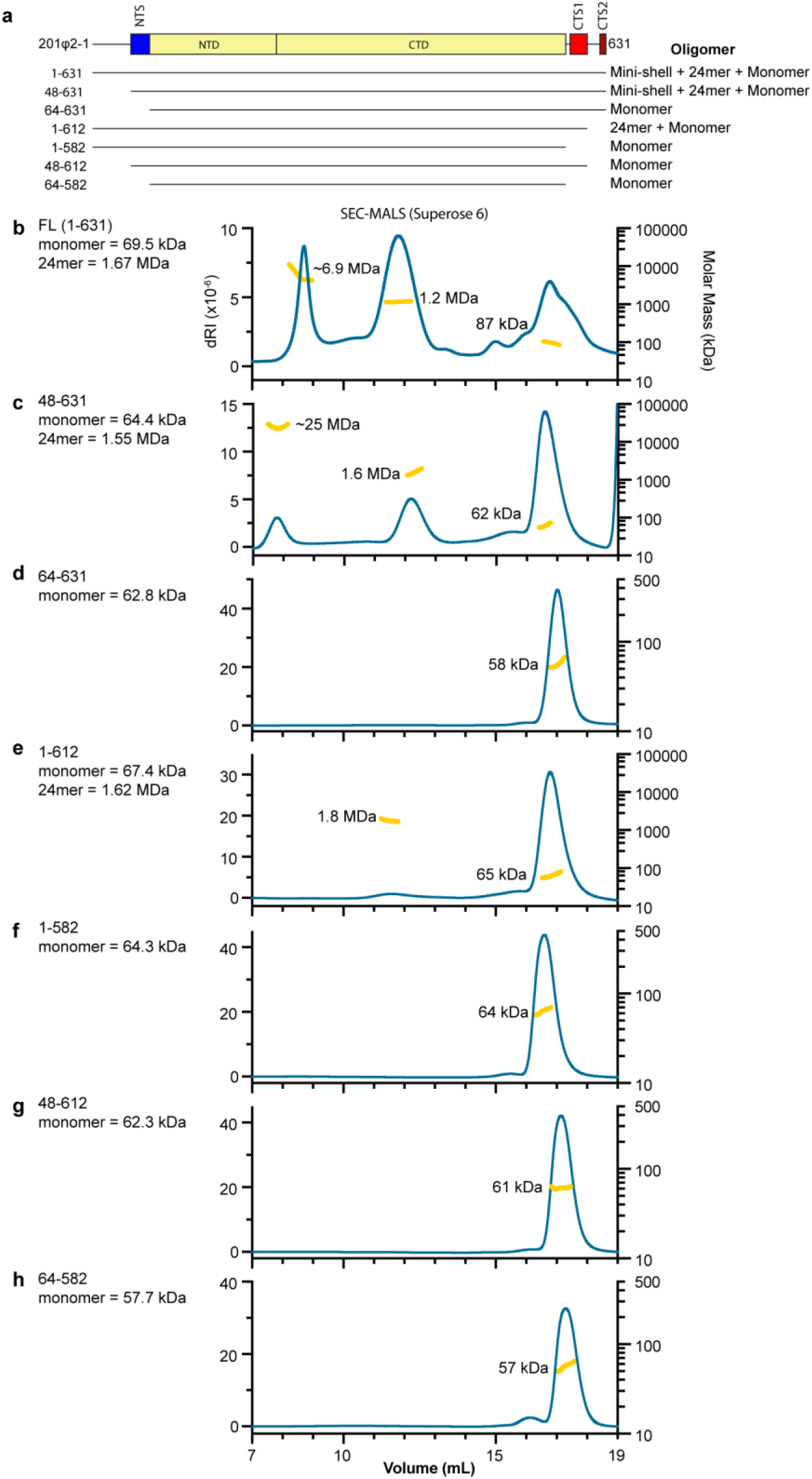
SEC-MALS of 201φ2-1 chimallinA truncations. **a**, Domain diagram of 201φ2-1 chimallinA (top), with truncations tested by SEC-MALS (bottom). **b-h**, SEC-MALS analysis of full-length 201φ2-1 chimallinA (**b**) and truncated constructs lacking the N-tail (**c**), NTS (**d**), CTS2 (**e**), CTS1+CTS2 (**f**), N-tail+CTS2 (**g**), or NTS + CTS1/2 (**h**). For panels b-h, differential refractive index (dRI) shows protein concentration (blue curves), and yellow points indicate measured molecular weight. Average molecular weight for each peak is shown.

**SI Figure 8.**
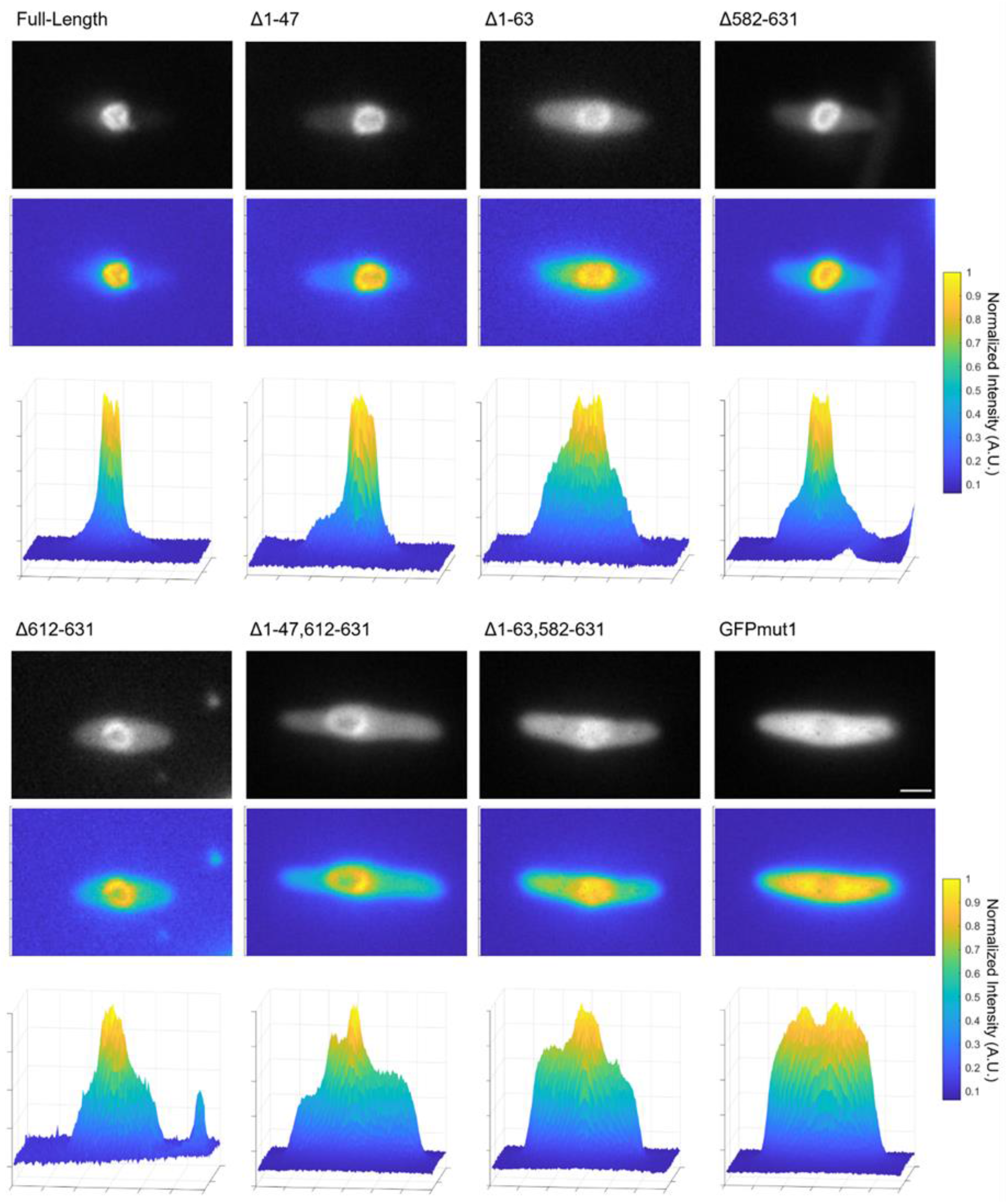
GFP-chimallinA incorporation into the nuclear shell in 201φ2-1-infected *P. chlororaphis*. Raw microscopy images of representative cells expressing GFP-chimallinA and infected with 201φ2-1 60 minutes post-infection (mpi) showing GFP fluorescence with associated 3D graphs showing normalized GFP fluorescence intensity within these cells from a top and side view. GFPmut1 was expressed without fusion to chimallinA as a negative control and shows no incorporation. Scale bar is 1 μm.

**SI Figure 9.**
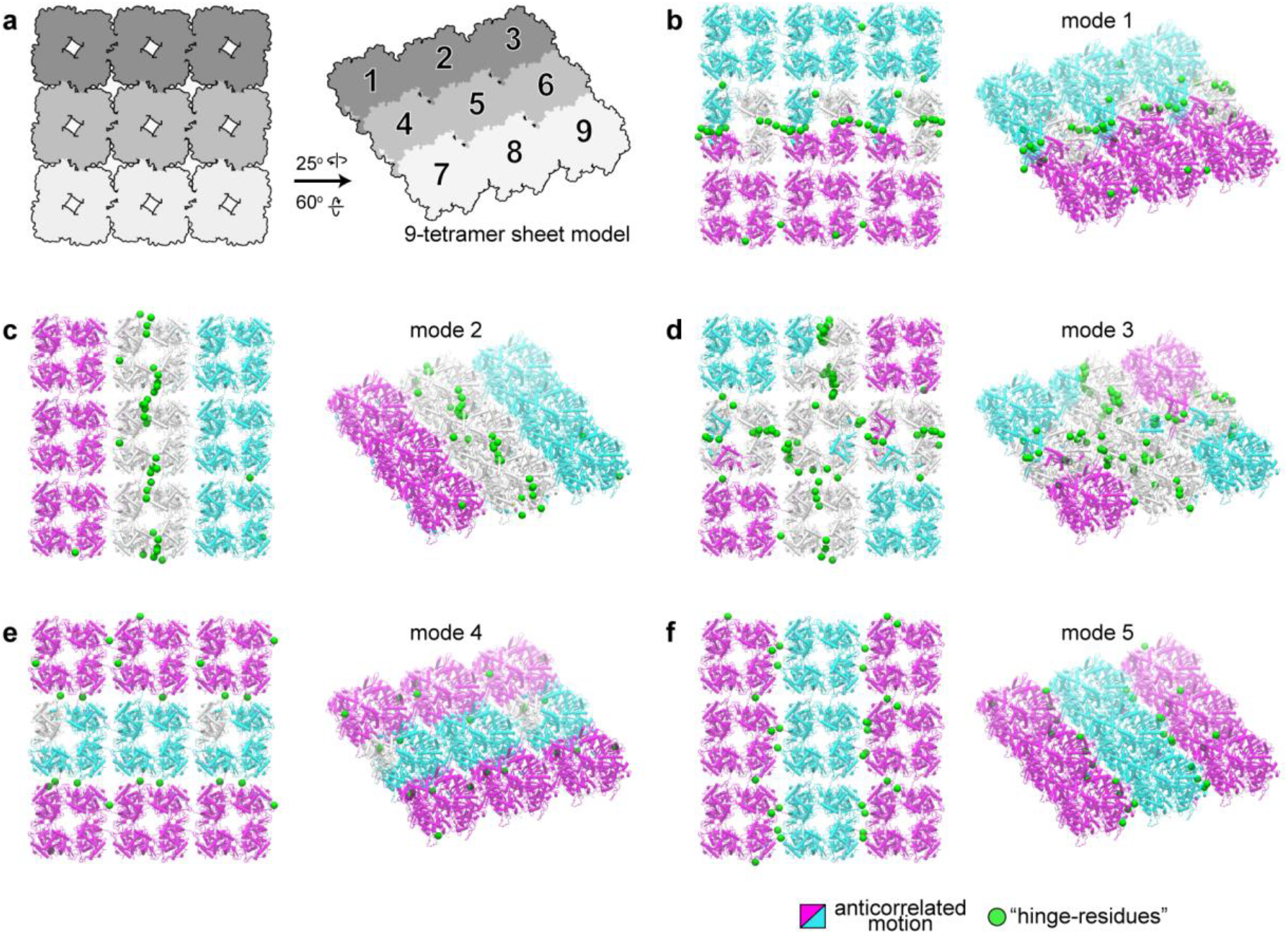
Gaussian network model analysis results. **a**, Cytosolic and tilted view schematic of 3×3 tetramer sheet model. **b-f**, Cartoon sheet model colored by results of Gaussian Network Model modes 1 through 5, respectively. Regions are colored according to the directional correlation of motion: positive (cyan), negative (magenta), and near-zero (white). “Hinge-residues” are depicted as green spheres. A list of the hinge-residues for each mode is in **SI Table 7**.

**SI Figure 10.**
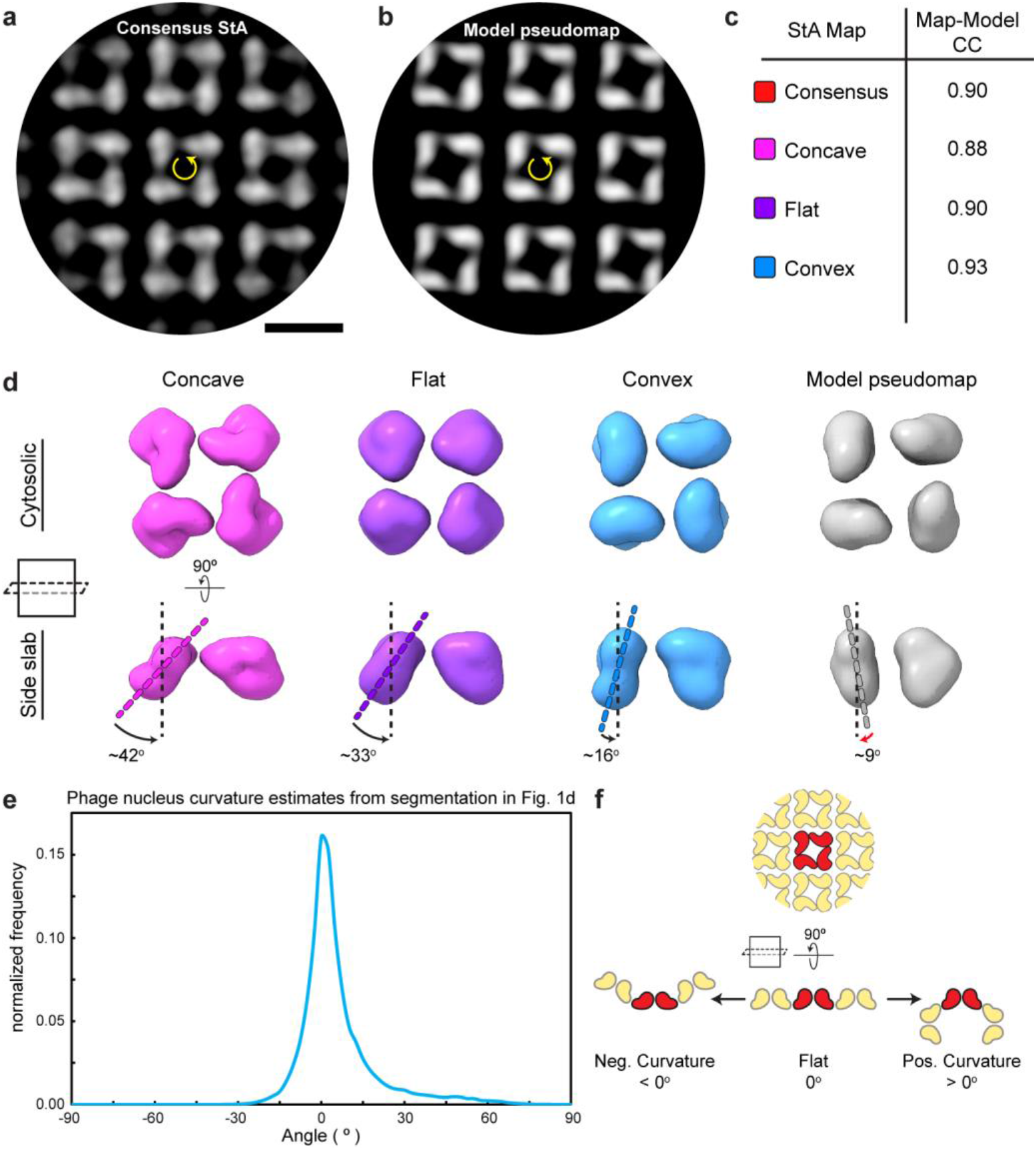
Comparison of 201φ2-1 single-particle and *in situ* subtomogram reconstructions. **a**, Slice through the consensus subtomogram average for the 201φ2-1 nuclear shell, with the four-fold axis defining the central tetramer noted. **b**, Equivalent view of panel **a** from a pseudomap generating by fitting tetramers into the consensus subtomogram average. **c**, Model-map correlation coefficient (CC) for a tetramer model fit into the consensus subtomogram average and the three subclasses (concave, flat, and convex). **d**, Two views of the concave, flat, and convex subclasses, compared to a pseudomap generated from the tetramer model. Dotted lines indicate the orientation of one monomer in each map. Denoted angles are with respect to the perpendicular. The arc arrow is red for the pseudomap to denote its opposite direction compared to the subtomogram average maps. **e**, Angles between tetramers in the phage nucleus lattice, derived from surface curvature estimates for phage nucleus segmentation in Figure 1d. **f**, Schematic of example manifestations of lattice curvature. The positive curvature shown on the right represents the 90° angle seen in the *in vitro* cubic assem-

**SI Figure 11.**
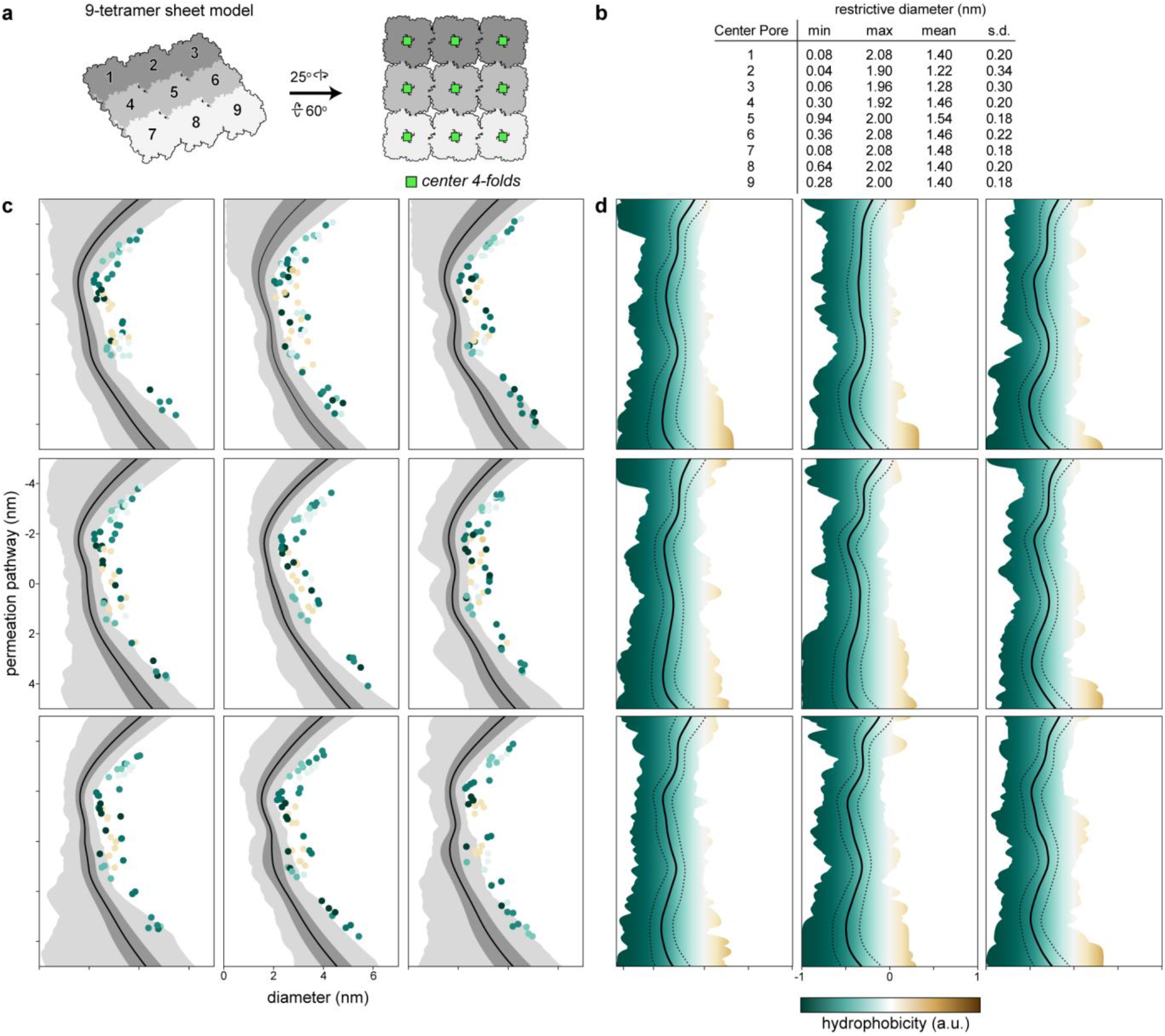
Size and hydrophobicity profiles of the center four-fold pores. **a**, Schematic of the 9-tetramer sheet model with the center four-fold pores marked with green squares. **b**, Pore diameter summary statistics for the nine center four-folds denoted in **a** over the course of the averaged 300-ns simulations (n = 5). **c**, Diameter profiles for each pore. The permeation pathway from top (negative values) to bottom (positive values) corresponds with cytosol to lumen. Solid black lines denote the mean diameter, dark gray shading +/- one standard deviation, and light grey shading the range. Dots indicate pore-facing residues and are colored by hydrophobicity. **d**, Hydrophobicity profiles for each pore. Solid black lines denote the mean hydrophobicity, dashed lines +/- one standard deviation, and shaded regions mark the range.

**SI Figure 12.**
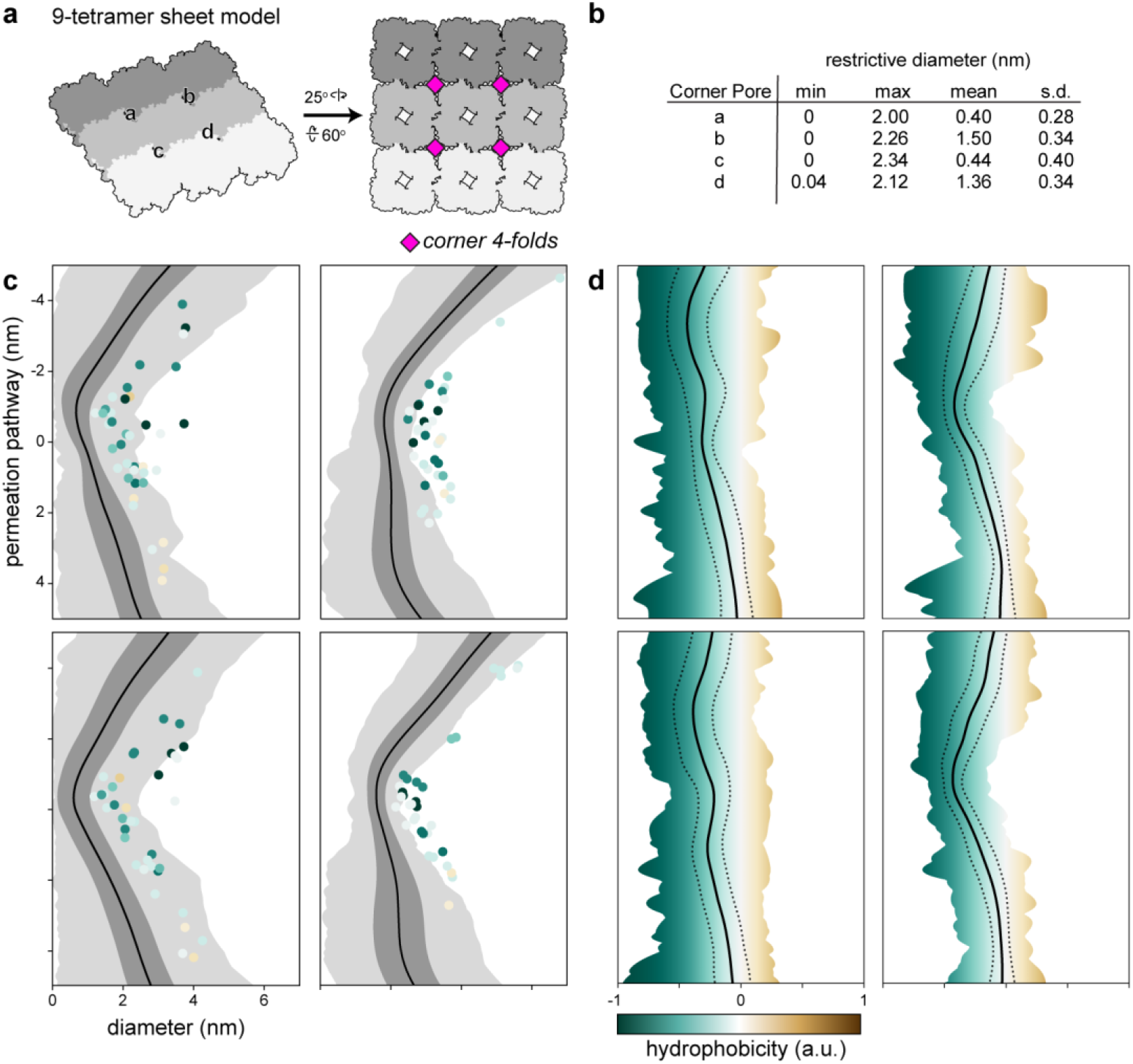
Size and hydrophobicity profiles of the corner four-fold pores. **a**, Schematic of the 9-tetramer sheet model with the corner four-fold pores marked with pink squares. **b**, Pore diameter summary statistics for the four corner four-folds denoted in **a** over the course of the averaged 300-ns simulations (n = 5). **c**, Diameter profiles for each pore. The permeation pathway from top (negative values) to bottom (positive values) corresponds with cytosol to lumen. Solid black lines denote the mean diameter, dark gray shading +/- one standard deviation, and light grey shading the range. Dots indicate pore-facing residues and are colored by hydrophobicity. **d**, Hydrophobicity profiles for each pore. Solid black lines denote the mean hydrophobicity, dashed lines +/- one standard deviation, and shaded regions mark the range.

**SI Figure 13.**
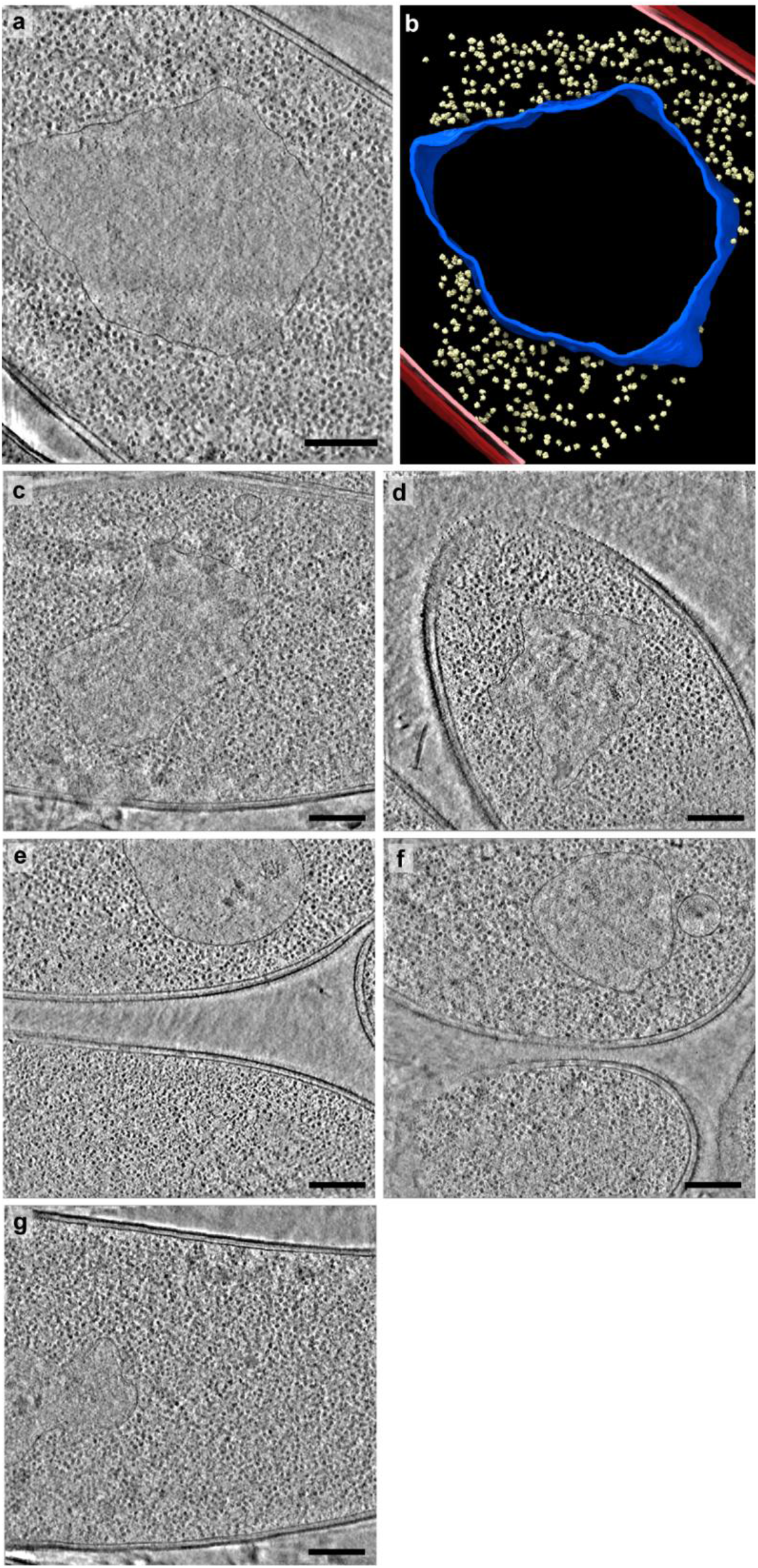
In situ cryoFIB-ET of Goslar infected APEC2248 cells. **a**, Tomographic slice of a Goslar-nucleus and **b** the corresponding segmentation model. Outer and inner bacterial membranes are burgundy and pink, respectively. The phage nucleus is colored blue and host ribosomes are colored pale yellow Five-hundred randomly selected 70S ribosomes are placed for clarity. (**c-g**) A slice from each of the Goslar-nucleus containing tomograms used for subtomogram averaging in this study. The cells were plunged at effectively ∼20-30 mpi, thus too early to observe virion assembly. All scale bars are 250 nm.

**SI Data Figure 14.**
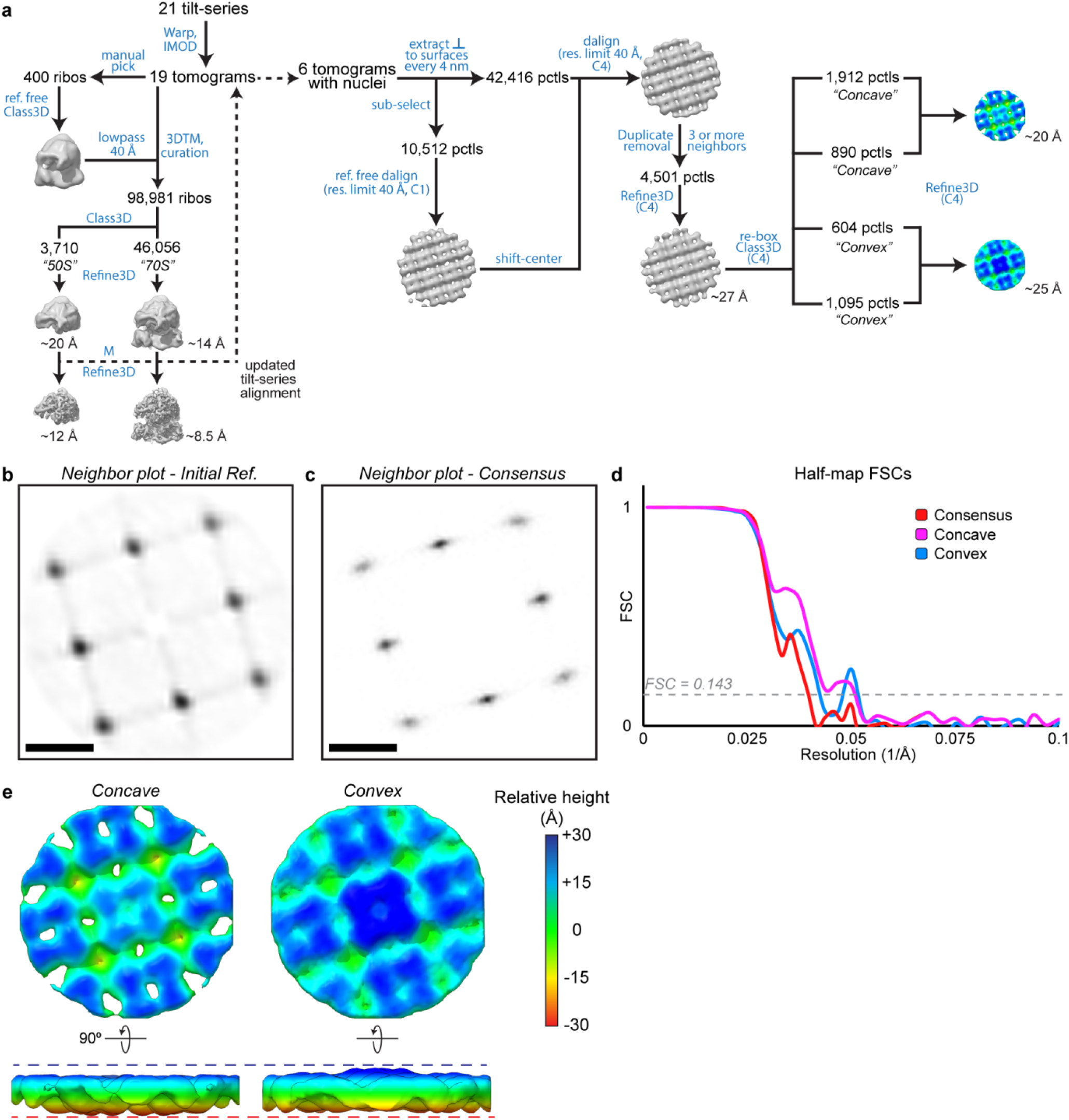
*In situ* subtomogram averaging of Goslar chimallinA in the context of the phage nucleus. **a**, Schematic of the subtomogram averaging workflow. **b**, Neighbor plot of the asymmetrically aligned initial reference. **c**, Neighbor plot of the symmetrized consensus refinement. **d**, Half-map FSC curves for the subtomogram reconstructions. **e**, Enlarged views of the resolved classes colored by relative height. Scale bar: 10 nm.

**SI Figure 15.**
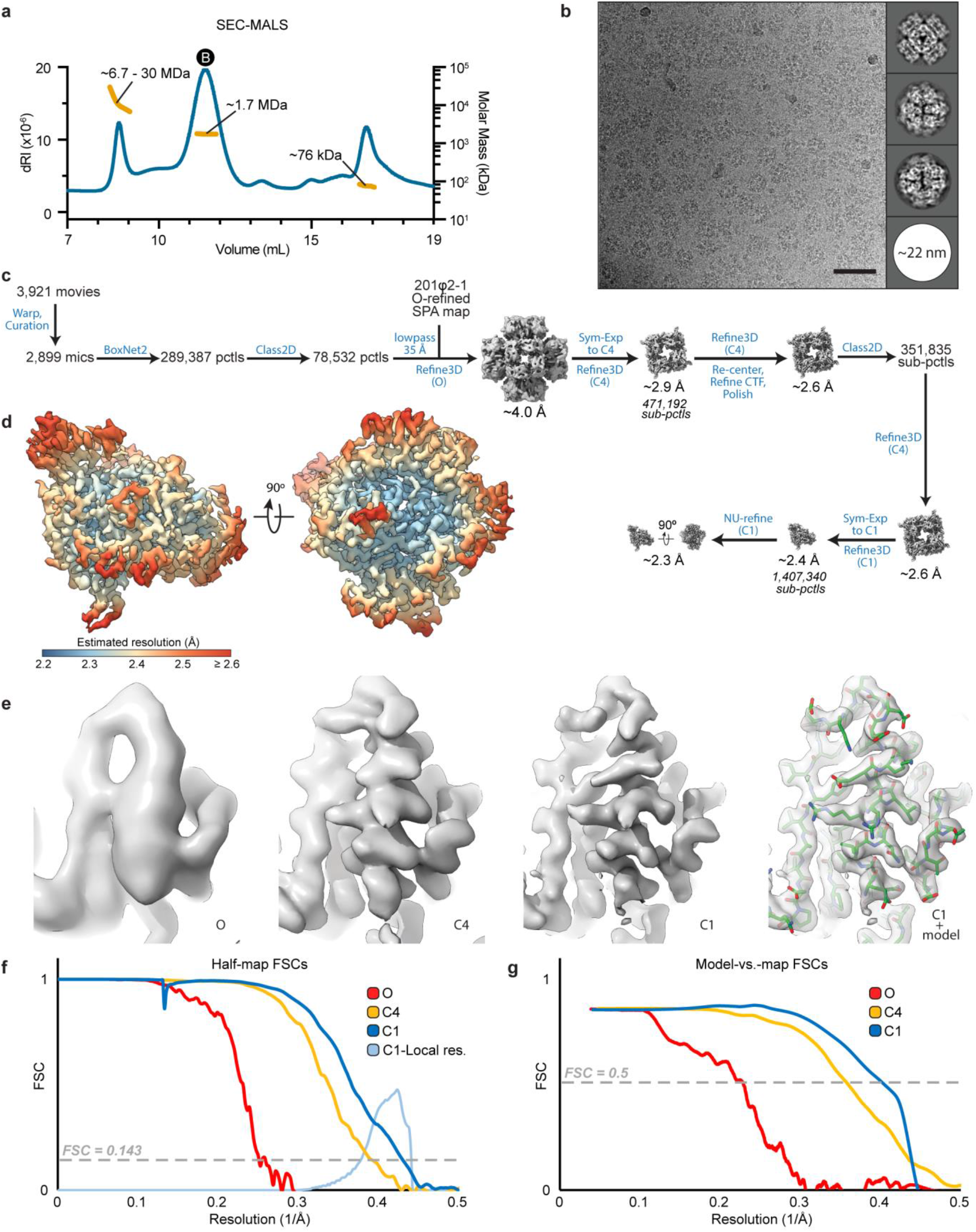
Single-particle reconstruction of the *in vitro* Goslar chimallinA cubic assembly. **a**, SEC-MALS analysis of purified, full-length Goslar chimallin. **b**, Exemplar micrograph and 2D class averages. Scale bar is 50 nm. **c**, Schematic of the localized reconstruction workflow. **d**, C1 reconstruction filtered and colored by local resolution estimate. **e**, Unsharpened density map views centered on helix B (residues 64-78) at progressive stages of the localized reconstruction process. Final view of the C1 map shown with fitted coordinate model. **f**,**g**, FSC curves for the half-maps and against corresponding models at progressive stages of the localized reconstruction process (red, yellow, and blue), histogram of local resolution estimates for the C1 reconstruction (light blue), and the C1 model-vs-map FSC curve (black).

**SI Figure 16.**
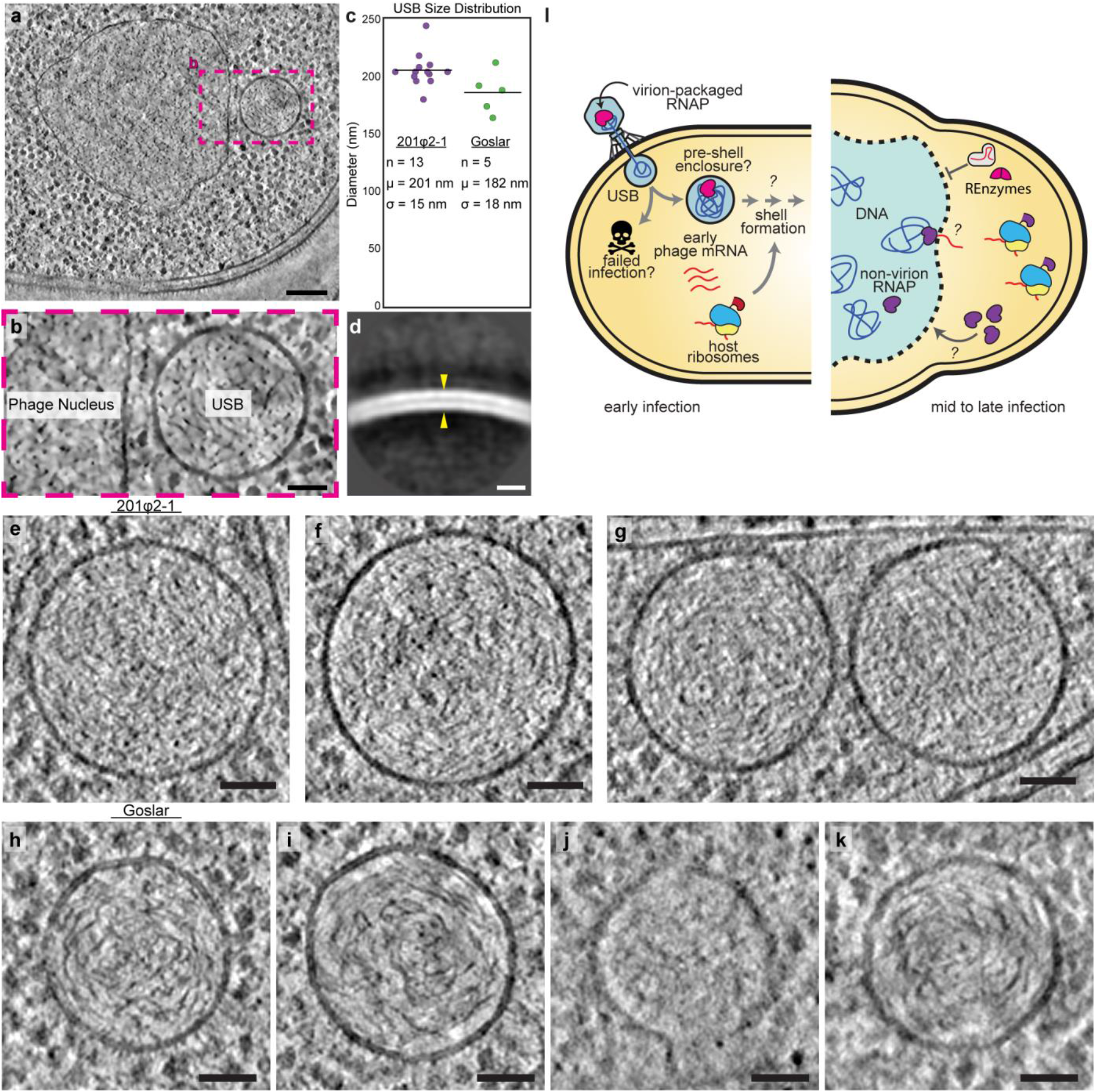
Unidentified spherical bodies present in jumbo phage-infected cell populations and speculative models. **a**, Tomographic slice of Goslar-infected APEC2248 cell containing a bonafide phage nucleus, as well as an unidentified spherical body (USB). **b**, Enlarged view of the phage nucleus and USB from the region boxed in **a. c**, Plot of the apparent maximal diameter distributions for 201φ2-1 (purple) and Goslar (green) USBs with the summary statistics listed. **d**, Subtomogram average of the USBs picked from the Goslar dataset. Yellow arrow pointing to putative membrane leaflets. Slices of USBs from the 201φ2-1 (**e-g**) and Goslar (**h-k**) datasets. **i**, Left, model of USBs as the previously proposed pre-shell/nucleus enclosure of the phage DNA (3). Right, schematic summary of structural models in this work: (i) exclusion of host nucleases by small chimallin pore sizes, (ii) possible extrusion of phage mRNA via these pores, and (iii) implication of additional shell components to enable uptake of specific phage proteins into the phage nucleus. Scale bars: **a**: 150 nm, **b**: 50 nm, **d**: 10 nm, **e-k**: 50 nm.

**SI Figure 17.**
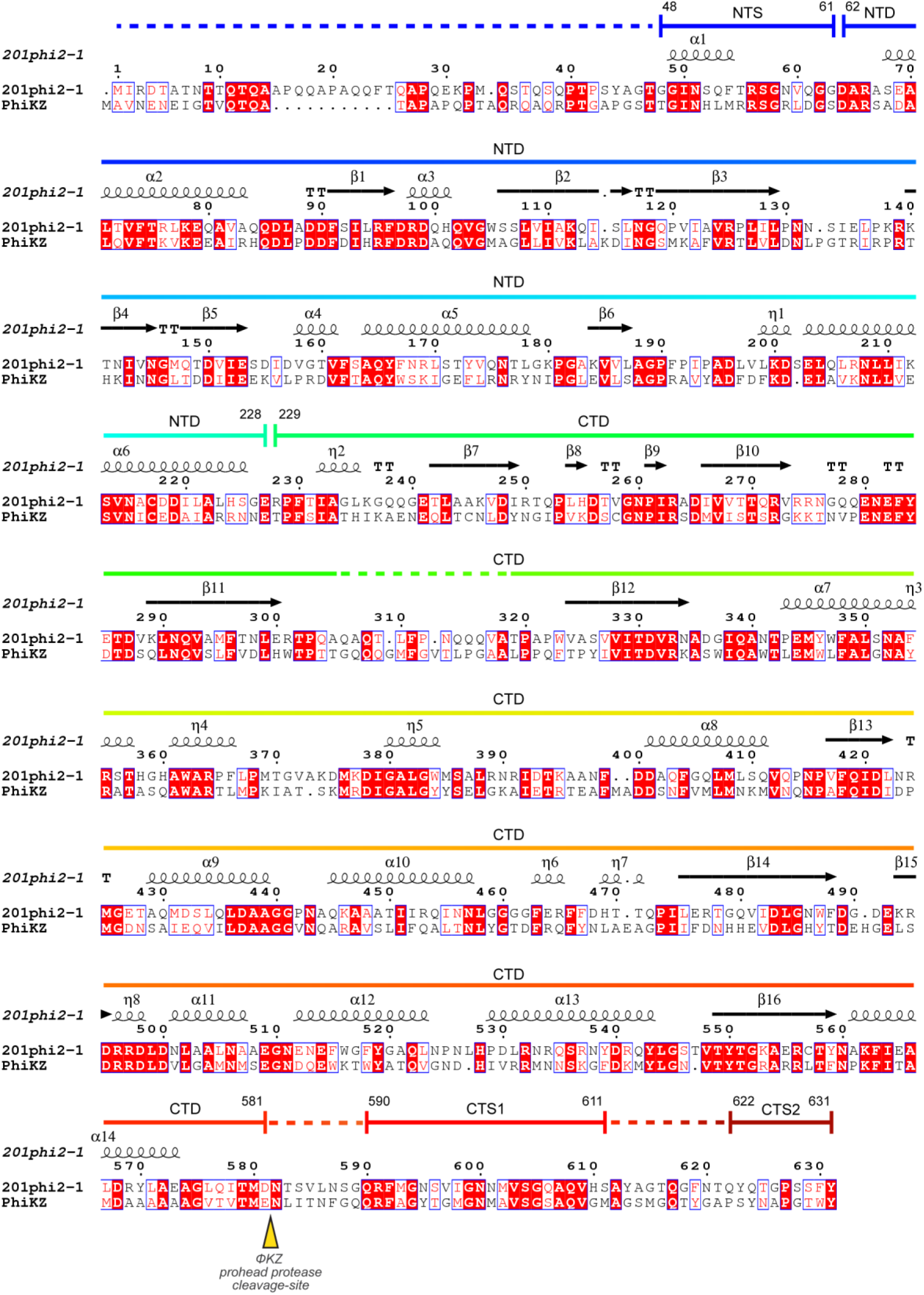
Alignment of 201φ2-1 and φKZ chimallinA proteins. Alignment of 201φ2-1 gp105 and φKZ gp54 protein sequences. Secondary structure and domain annotation are shown above the alignment and are based on the 201φ2-1 structure. Dashed lines in the domain annotation indicate unresolved segments. The φKZ prohead protease (gp175) cleavage-site in gp54 is indicated by a yellow arrow. Cleavage-site based on that identified by Weintraub *et al*. (20).

**SI Figure 18.**
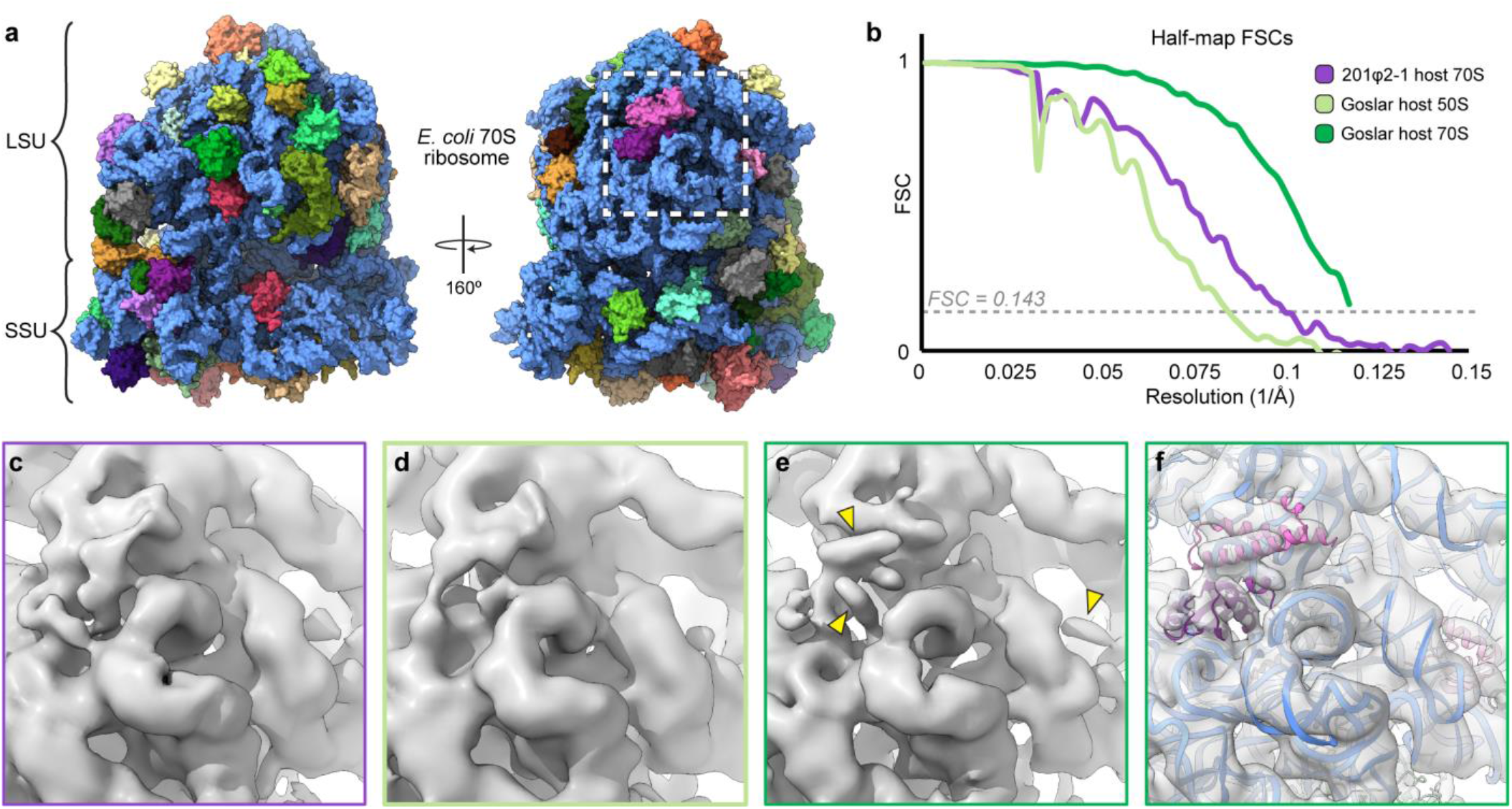
Subtomogram reconstructions of jumbo phage host ribosomes. **a**, Surface depictions of an *E. coli* 70S ribosome coordinate model (PDB ID: 7K00) with rRNA in pale blue and protein subunits in a variety of colors. **b**, Half-map FSC curves for the jumbo phage host ribosome reconstructions obtained in this study. **c-f**, Close-ups of the region boxed in **a** showing the density of the various reconstructions. Panel outline color matches the scheme in **b. c**, 201φ2-1 host (*P. chlororaphis*) 70S. **d**, Goslar host (APEC 2248) 50S. **e**, Goslar host (APEC 2248) 70S with yellow arrows pointing to density features corresponding to alpha-helices. **f**, Goslar host (APEC 2248) 70S with a cartoon depiction of the coordinate model from **a** docked into the map. The 23S rRNA is pale blue, L29 is pink, L23 is dark purple, and L28 is light pink.

**SI Table 1.** *In situ* cryoFIB-ET data collection and reconstruction statistics.

|  | 201φ2-1 infected *P. chlororaphis* |  |  |  |  | Goslar infected *E. coli* (APEC2248) |  |  |  |  |
| --- | --- | --- | --- | --- | --- | --- | --- | --- | --- | --- |
|  | chimallinA |  |  |  |  | chimallinA |  |  |  |  |
|  | 70S<br>(EMD-25183) | Consensus<br>(EMD-25221) | Concave<br>(EMD-25220) | Flat<br>(EMD-25222) | Convex<br>(EMD-25223) | 50S<br>(EMD-25360) | 70S<br>(EMD-25359) | Consensus<br>(EMD-25229) | Concave<br>(EMD-25262) | Convex<br>(EMD-25358) |
| **Data collection** |  |  |  |  |  |  |  |  |  |  |
| Magnification |  |  | 33,000 |  |  |  |  | 42,000 |  |  |
| Voltage (kV) |  |  |  |  |  | 300 |  |  |  |  |
| Defocus range<br>(μm) |  |  | -4.5 to -6 |  |  |  |  | -5 to -6 |  |  |
| Pixel size (Å) |  |  | 3.46 |  |  |  |  | 4.27 |  |  |
| Electron exposure<br>(e-/Å<sup>2</sup>) |  |  | 120 |  |  |  |  | 150 |  |  |
| Tilt-range/step (°) |  |  | ±51 / 3 |  |  |  |  | ±56 / 2 |  |  |
| Tilt-scheme |  |  |  |  | dose-symmetric, grouping 2 |  |  |  |  |  |
| **Processing** |  |  |  |  |  |  |  |  |  |  |
| Symmetry imposed | C1 | C4 | C4 | C4 | C4 | C1 | C1 | C4 | C4 | C4 |
| Initial particle im-<br>ages (no.) | 17,169 | 66,887 |  | 8,454 |  | 98,981 |  | 42,416 | 4,501 |  |
| Final particle im-<br>ages (no.) | 11,148 | 8,454 | 2,033 | 4,475 | 945 | 3,710 | 46,056 | 4,501 | 2,802 | 1,699 |
| Map resolution (Å) | 10 | 24 | 20 | 18 | 23 | 12 | 8.5 | 27 | 20 | 25 |
| FSC threshold |  |  |  |  | 0.143 |  |  |  |  |  |

**SI Table 2.** Cryo-EM data collection, reconstruction, and refinement statistics.

|  | 201φ2-1<br>gp105 |  |  | Goslar<br>gp189 |  |  |
| --- | --- | --- | --- | --- | --- | --- |
|  | 24mer<br>(EMD-25390)<br>(PDB 7SQQ) | tetramer<br>(EMD-25391)<br>(PDB 7SQR) | monomer<br>(EMD-25392)<br>(PDB 7SQS) | 24mer<br>(EMD-25394)<br>(PDB 7SQT) | tetramer<br>(EMD-25395)<br>(PDB 7SQU) | monomer<br>(EMD-25396)<br>(PDB 7SQV) |
| **Data collection and processing** |  |  |  |  |  |  |
| Magnification |  | 130,000 |  |  | 165,000 |  |
| Voltage (kV) |  |  | 300 |  |  |  |
| Electron exposure (e-/Å<sup>2</sup>) |  | 43 |  |  | 40 |  |
| Defocus range (μm) |  |  | -0.1 to -1.5 |  |  |  |
| Pixel size (Å) |  | 1.075 |  |  | 0.8452 |  |
| Symmetry imposed | O | C4 | C1 | O | C4 | C1 |
| Initial particle images (no.) | 140,782 | 772,788<sup>a</sup> | 2,657,452<sup>a</sup> | 289,387 | 471,192<sup>a</sup> | 1,407,340<sup>a</sup> |
| Final particle images (no.) | 128,798 | 664,363<sup>a</sup> | 2,657,452<sup>a</sup> | 78,532 | 351,835<sup>a</sup> | 1,407,340<sup>a</sup> |
| Map resolution (Å) | 4.2 | 3.4 | 3.1 | 4.0 | 2.6 | 2.3 |
| FSC threshold |  |  | 0.143 |  |  |  |
| Map resolution range (Å) | 4.0 to 5.1 | 3.3 to 4.4 | 2.9 to 4.8 | 3.7 to 6.0 | 2.5 to 3.4 | 2.3 to 3.4 |
| **Refinement** |  |  |  |  |  |  |
| Initial model used (PDB code) |  | *de novo* |  |  | 7SQS |  |
| Model resolution (Å) | 4.7 | 3.6 | 3.3 | 4.4 | 2.8 | 2.5 |
| FSC threshold |  |  | 0.5 |  |  |  |
| Model composition |  |  |  |  |  |  |
| Non-hydrogen atoms | 103,632 | 17,272 | 4,376 | 107,136 | 17,856 | 4,471 |
| Protein residues | 13,296 | 2,216 | 560 | 13,752 | 2,292 | 573 |
| Ligands | 0 | 0 | 0 | 0 | 0 | 7 |
| B factors (Å<sup>2</sup>) |  |  |  |  |  |  |
| Protein | 408.17 | 125.69 | 108.00 | 318.81 | 97.77 | 77.87 |
| Ligand (waters) | n.a. | n.a. | n.a. | n.a. | n.a. | 71.61 |
| R.m.s. deviations |  |  |  |  |  |  |
| Bond lengths (Å) | 0.003 | 0.003 | 0.002 | 0.010 | 0.010 | 0.001 |
| Bond angles (°) | 0.633 | 0.632 | 0.469 | 0.699 | 0.698 | 0.380 |
| Validation |  |  |  |  |  |  |
| MolProbity score | 1.63 | 1.56 | 1.31 | 1.16 | 0.94 | 0.87 |
| Clashscore | 8.23 | 6.82 | 3.59 | 2.13 | 1.16 | 1.36 |
| Poor rotamers (%) | 0 | 0 | 0.22 | 1.25 | 1.04 | 0.63 |
| Ramachandran plot |  |  |  |  |  |  |
| Favored (%) | 96.89 | 96.89 | 97.08 | 97.53 | 97.53 | 99.29 |
| Allowed (%) | 3.11 | 3.11 | 2.92 | 2.47 | 2.47 | 0.71 |
| Disallowed (%) | 0 | 0 | 0 | 0 | 0 | 0 |

<sup>a</sup>Denotes count of sub-particle images

**SI Table 3.**
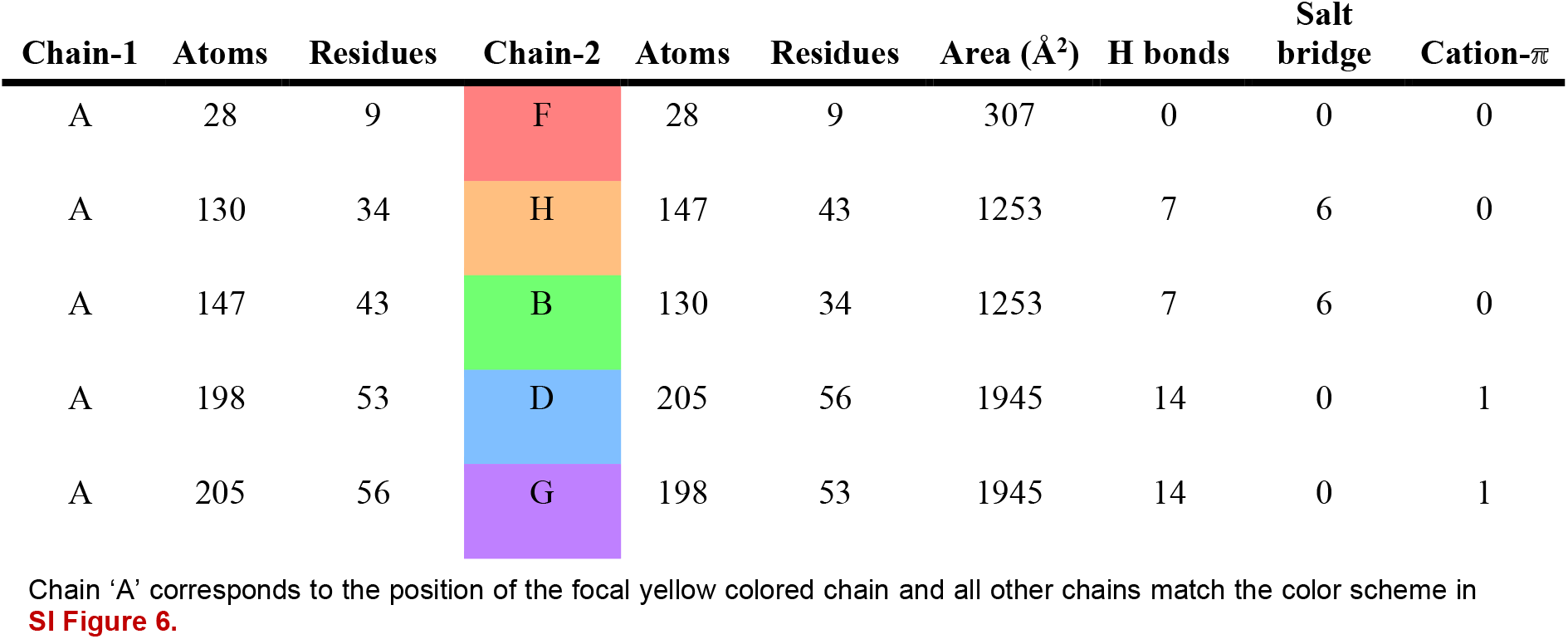
Summary of *in vitro* 201φ2-1 chimallinA protomer interfaces.

| Chain-1 | Atoms | Residues | Chain-2 | Atoms | Residues | Area (Å<sup>2</sup>) | H bonds | Salt bridge | Cation-π |
| --- | --- | --- | --- | --- | --- | --- | --- | --- | --- |
| A | 28 | 9 | F | 28 | 9 | 307 | 0 | 0 | 0 |
| A | 130 | 34 | H | 147 | 43 | 1253 | 7 | 6 | 0 |
| A | 147 | 43 | B | 130 | 34 | 1253 | 7 | 6 | 0 |
| A | 198 | 53 | D | 205 | 56 | 1945 | 14 | 0 | 1 |
| A | 205 | 56 | G | 198 | 53 | 1945 | 14 | 0 | 1 |

Chain 'A' corresponds to the position of the focal yellow colored chain and all other chains match the color scheme in **SI Figure 6**.

**SI Table 4.** Summary of a 201φ2-1 chimallinA protomer’s polar interactions *in vitro*.

| Chain-1 |  | Resi | Resn | Atom | Chain-2 |  | Resi | Resn | Atom | Dist (Å) | Type |
| --- | --- | --- | --- | --- | --- | --- | --- | --- | --- | --- | --- |
| A | B | 48 | GLY | N | H | A | 287 | ASP | OD2 | 3.6 | H bond |
| A | B | 51 | ASN | NE2 | H | A | 337 | GLY | O | 3.2 | H bond |
| A | B | 51 | ASN | OD1 | H | A | 339 | GLN | N | 3.4 | H bond |
| A | B | 56 | ARG | O | H | A | 339 | GLN | NE2 | 2.6 | H bond |
| A | B | 59 | ASN | N | H | A | 86 | ASP | O | 3.5 | H bond |
| A | B | 102 | GLN | OE1 | H | A | 281 | ASN | ND2 | 2.3 | H bond |
| A | B | 163 | SER | OG | H | A | 282 | GLU | OE2 | 3.1 | H bond |
| A | B | 56 | ARG | NE | H | A | 219 | ASP | OD1 | 2.5 | Salt bridge |
| A | B | 56 | ARG | NH1 | H | A | 227 | GLU | OE1 | 3.4 | Salt bridge |
| A | B | 56 | ARG | NH1 | H | A | 227 | GLU | OE2 | 3.7 | Salt bridge |
| A | B | 56 | ARG | NH2 | H | A | 227 | GLU | OE1 | 3.9 | Salt bridge |
| A | B | 56 | ARG | NH2 | H | A | 227 | GLU | OE2 | 2.9 | Salt bridge |
| A | B | 56 | ARG | NH2 | H | A | 219 | ASP | OD1 | 3.3 | Salt bridge |
| A | G | 319 | THR | OG1 | D | A | 319 | THR | N | 2.9 | H bond |
| A | G | 406 | GLN | OE1 | D | A | 306 | GLN | N | 3.9 | H bond |
| A | G | 424 | ARG | NH2 | D | A | 624 | GLN | O | 3.3 | H bond |
| A | G | 430 | GLN | OE1 | D | A | 629 | SER | N | 3.8 | H bond |
| A | G | 430 | GLN | OE1 | D | A | 629 | SER | OG | 2.8 | H bond |
| A | G | 437 | ASP | OD2 | D | A | 631 | TYR | OH | 2.8 | H bond |
| A | G | 592 | PHE | O | D | A | 516 | TRP | NE1 | 2.3 | H bond |
| A | G | 595 | ASN | N | D | A | 301 | ARG | O | 3.5 | H bond |
| A | G | 604 | SER | N | D | A | 565 | GLU | OE2 | 3 | H bond |
| A | G | 604 | SER | OG | D | A | 565 | GLU | OE2 | 3.4 | H bond |
| A | G | 605 | GLY | N | D | A | 565 | GLU | OE2 | 3.6 | H bond |
| A | G | 606 | GLN | NE2 | D | A | 358 | HIS | O | 3.1 | H bond |
| A | G | 607 | ALA | N | D | A | 357 | THR | OG1 | 3.8 | H bond |
| A | G | 610 | HIS | N | D | A | 578 | ILE | O | 3.1 | H bond |

Chain 'A' corresponds to the position of the focal yellow colored chain and all other chains match the color scheme in **SI Figure 6**.

**SI Table 5.** Summary of *in vitro* Goslar chimallinA protomer interfaces.

| Chain-1 | Atoms | Residues | Chain-2 | Atoms | Residues | Area (Å<sup>2</sup>) | H bonds | Salt<br>bridge | Cation-π |
| --- | --- | --- | --- | --- | --- | --- | --- | --- | --- |
| A | 11 | 3 | F | 11 | 3 | 91 | 0 | 2 | 0 |
| A | 164 | 38 | H | 210 | 62 | 1779 | 18 | 11 | 1 |
| A | 210 | 62 | B | 164 | 38 | 1779 | 18 | 11 | 1 |
| A | 206 | 54 | D | 210 | 58 | 2024 | 16 | 6 | 3 |
| A | 210 | 58 | G | 206 | 56 | 2024 | 16 | 6 | 3 |

Chain 'A' corresponds to the position of the focal yellow colored chain and all other chains match the color scheme in **SI Figure 6**.

**SI Table 6.** Summary of a Goslar chimallinA protomer’s polar interactions *in vitro*.

| Chain-1 |  | Resi | Resn | Atom | Chain-2 |  | Resi | Resn | Atom | Dist (Å) | Type |
| --- | --- | --- | --- | --- | --- | --- | --- | --- | --- | --- | --- |
| A | F | 140 | HIS | NE2 | F | A | 427 | ASP | OD2 | 3.2 | Salt bridge |
| A | B | 46 | MET | O | H | A | 279 | GLN | NE2 | 3.5 | H bond |
| A | B | 50 | ASN | ND2 | H | A | 330 | GLU | O | 3.4 | H bond |
| A | B | 50 | ASN | OD1 | H | A | 332 | ASP | N | 3.5 | H bond |
| A | B | 55 | ARG | NE | H | A | 206 | ASN | OD1 | 3.1 | H bond |
| A | B | 55 | ARG | NH2 | H | A | 210 | THR | OG1 | 3.2 | H bond |
| A | B | 56 | ARG | NH1 | H | A | 107 | SER | OG | 3.7 | H bond |
| A | B | 56 | ARG | NH2 | H | A | 209 | GLN | OE1 | 3.4 | H bond |
| A | B | 56 | ARG | NH2 | H | A | 107 | SER | OG | 3.6 | H bond |
| A | B | 57 | ILE | N | H | A | 205 | ASP | OD2 | 3.1 | H bond |
| A | B | 57 | ILE | O | H | A | 202 | SER | OG | 3.6 | H bond |
| A | B | 59 | ARG | NH1 | H | A | 332 | ASP | O | 3.7 | H bond |
| A | B | 59 | ARG | NH2 | H | A | 331 | LEU | O | 3.1 | H bond |
| A | B | 60 | ASN | ND1 | H | A | 329 | LEU | O | 3.3 | H bond |
| A | B | 97 | SER | O | H | A | 275 | TYR | OH | 3.2 | H bond |
| A | B | 130 | PRO | O | H | A | 278 | GLN | NE2 | 2.7 | H bond |
| A | B | 152 | ASN | ND2 | H | A | 276 | ASP | O | 3.3 | H bond |
| A | B | 152 | ASN | OD1 | H | A | 276 | ASP | N | 3.3 | H bond |
| A | B | 154 | ASN | ND2 | H | A | 273 | SER | O | 3.3 | H bond |
| A | B | 55 | ARG | NH1 | H | A | 213 | GLU | OE1 | 3.8 | Salt bridge |
| A | B | 55 | ARG | NH1 | H | A | 213 | GLU | OE2 | 3.1 | Salt bridge |
| A | B | 55 | ARG | NH2 | H | A | 213 | GLU | OE1 | 3.4 | Salt bridge |
| A | B | 56 | ARG | NE | H | A | 205 | ASP | OD1 | 3.1 | Salt bridge |
| A | B | 56 | ARG | NH1 | H | A | 85 | ASP | OD2 | 3.3 | Salt bridge |
| A | B | 56 | ARG | NH2 | H | A | 205 | ASP | OD1 | 3.1 | Salt bridge |
| A | B | 134 | ARG | NE | H | A | 276 | ASP | OD2 | 3.9 | Salt bridge |
| A | B | 134 | ARG | NH2 | H | A | 274 | ASP | OD1 | 3.5 | Salt bridge |
| A | B | 134 | ARG | NH2 | H | A | 274 | ASP | OD1 | 3.5 | Salt bridge |
| A | B | 134 | ARG | NH2 | H | A | 274 | ASP | OD2 | 3.2 | Salt bridge |
| A | B | 134 | ARG | NH2 | H | A | 276 | ASP | OD2 | 3.6 | Salt bridge |
| A | B | 147 | ARG | NH1 | H | A | 276 | ASP | OD2 | 3.6 | Salt bridge |
| A | B | 56 | ARG | CZ | H | A | 86 | PHE | - | 5.6 | Cation-π |
| A | G | 399 | TYR | OH | D | A | 300 | ASN | N | 3.5 | H bond |
| A | G | 403 | GLN | NE2 | D | A | 297 | PRO | O | 3.4 | H bond |
| A | G | 427 | ARG | NH2 | D | A | 628 | PHE | O | 2.5 | H bond |
| A | G | 593 | ARG | NH2 | D | A | 517 | GLU | O | 4.7 | H bond |
| A | G | 595 | ASN | N | D | A | 292 | TYR | O | 2.9 | H bond |
| A | G | 595 | ASN | O | D | A | 293 | SER | OG | 2.9 | H bond |
| A | G | 599 | ARG | NE | D | A | 293 | SER | OG | 3.6 | H bond |
| A | G | 599 | ARG | NH2 | D | A | 294 | PRO | O | 3.3 | H bond |
| A | G | 599 | ARG | NH2 | D | A | 293 | SER | OG | 3.7 | H bond |
| A | G | 603 | LEU | N | D | A | 410 | LEU | O | 2.7 | H bond |
| A | G | 603 | LEU | O | D | A | 410 | LEU | N | 3.6 | H bond |
| A | G | 604 | GLY | O | D | A | 351 | ASN | ND2 | 3.5 | H bond |
| A | G | 605 | ASN | O | D | A | 566 | GLN | NE2 | 2.8 | H bond |
| A | G | 610 | ILE | O | D | A | 579 | ILE | N | 2.9 | H bond |
| A | G | 611 | TYR | OH | D | A | 348 | ALA | O | 3.2 | H bond |
| A | G | 612 | ALA | N | D | A | 579 | ILE | O | 3.4 | H bond |
| A | G | 593 | ARG | NE | D | A | 521 | ASP | OD1 | 3.0 | Salt bridge |
| A | G | 593 | ARG | NH1 | D | A | 520 | GLU | OE1 | 3.5 | Salt bridge |
| A | G | 593 | ARG | NH2 | D | A | 521 | ASP | OD1 | 3.6 | Salt bridge |
| A | G | 593 | ARG | NH2 | D | A | 520 | GLU | OE1 | 2.6 | Salt bridge |
| A | G | 601 | ARG | NH1 | D | A | 559 | ASP | OD1 | 3.4 | Salt bridge |
| A | G | 601 | ARG | NH1 | D | A | 559 | ASP | OD2 | 3.1 | Salt bridge |
| A | G | 427 | ARG | CZ | D | A | 631 | PHE | - | 4.3 | Cation-π |
| A | G | 593 | ARG | CZ | D | A | 292 | TYR | - | 5.2 | Cation-π |

Chain 'A' corresponds to the position of the focal yellow colored chain and all other chains match the color scheme in SI Figure 6.

**SI Figure 7.**
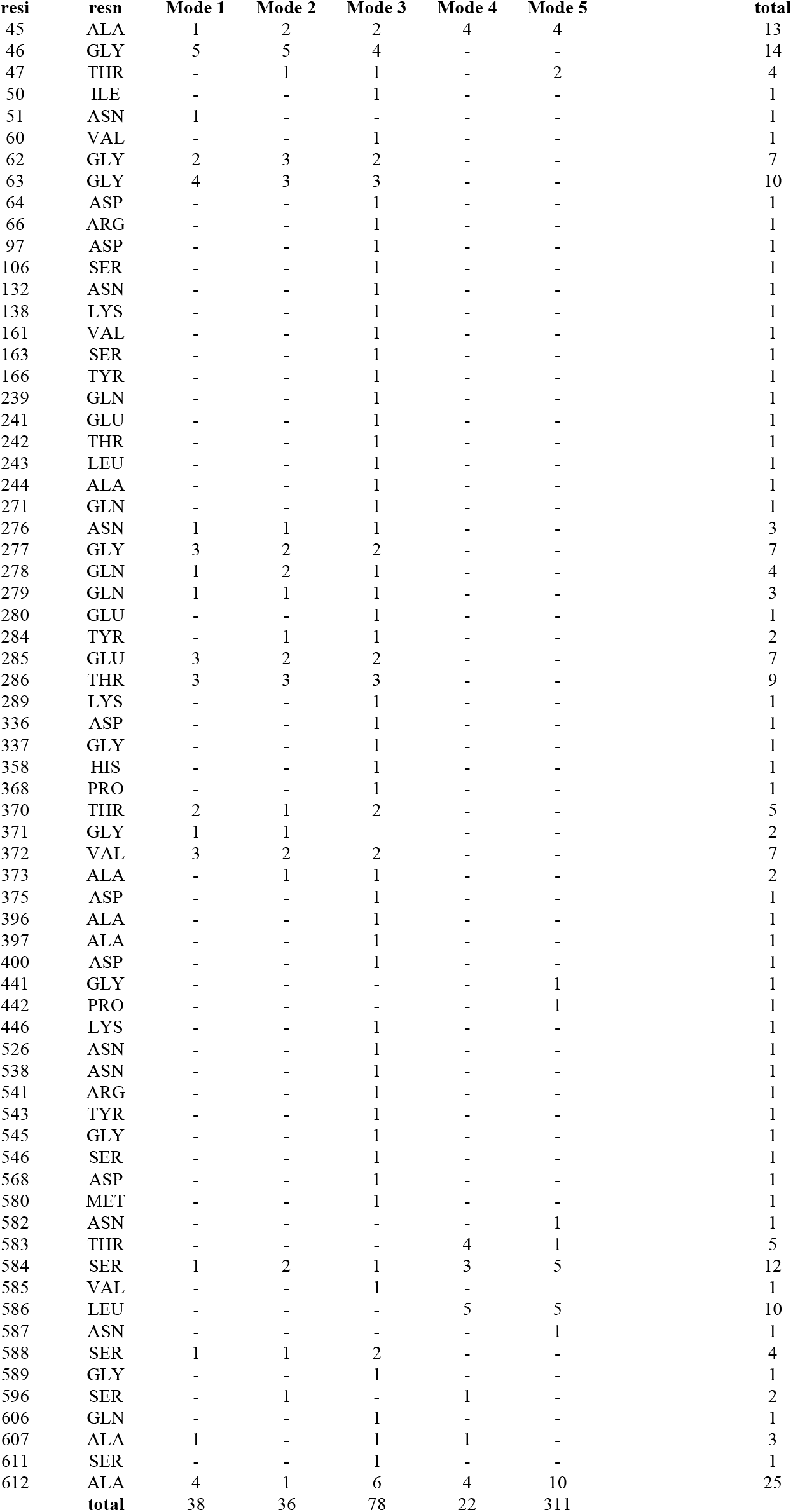
Gaussian network model hinge residues for 201φ2-1 chimallinA.

**SI Table 8.** Pore-facing residues^a^ during 201φ2-1 chimallinA simulations.

| Center 4-fold |  |  | Corner 4-fold |  |  |
| --- | --- | --- | --- | --- | --- |
| resi | resn | count | resi | resn | count |
| 69 | GLU | 2 | 117 | ASN | 3 |
| 72 | THR | 6 | 119 | GLN | 2 |
| 76 | ARG | 9 | 303 | PRO | 1 |
| 79 | GLU | 9 | 304 | GLN | 1 |
| 80 | GLN | 6 | 305 | ALA | 1 |
| 83 | ALA | 5 | 307 | ALA | 1 |
| 131 | ASN | 1 | 309 | THR | 1 |
| 133 | SER | 1 | 311 | PHE | 1 |
| 196 | ASP | 2 | 312 | PRO | 1 |
| 198 | VAL | 9 | 317 | VAL | 1 |
| 200 | LYS | 9 | 319 | THR | 1 |
| 201 | ASP | 8 | 320 | PRO | 2 |
| 203 | ARG | 1 | 414 | PRO | 1 |
| 203 | GLU | 3 | 415 | ASN | 1 |
| 372 | VAL | 4 | 445 | GLN | 1 |
| 373 | ALA | 1 | 452 | ILE | 1 |
| 374 | LYS | 8 | 453 | ARG | 2 |
| 375 | ASP | 2 | 456 | ASN | 1 |
| 377 | LYS | 1 | 464 | GLU | 4 |
| 378 | ASP | 1 | 465 | ARG | 2 |
| 391 | ARG | 5 | 466 | PHE | 1 |
| 395 | LYS | 4 | 468 | ASP | 3 |
| 480 | GLY | 1 | 469 | HIS | 1 |
| 488 | PHE | 9 | 470 | THR | 4 |
| 490 | GLY | 8 | 471 | THR | 4 |
| 491 | ASP | 4 | 472 | GLN | 2 |
| 493 | LYS | 6 | 562 | LYS | 2 |
| 509 | ASP | 1 | 569 | ARG | 1 |
| 509 | GLU | 8 | 573 | GLU | 1 |
| 511 | ASN | 1 | 596 | SER | 1 |
| 546 | SER | 5 | 600 | ASN | 3 |
| 547 | THR | 5 | 601 | ASN | 1 |
|  |  |  | 602 | MET | 1 |
|  |  |  | 604 | SER | 2 |
|  |  |  | 606 | GLN | 1 |
|  |  |  | 607 | ALA | 1 |

<sup>a</sup>Pore-facing residues are defined as residues exposing their amino acid side chains to the channel pore more than 50% over the course of the averaged 300-ns simulations.

**SI Table 9.**
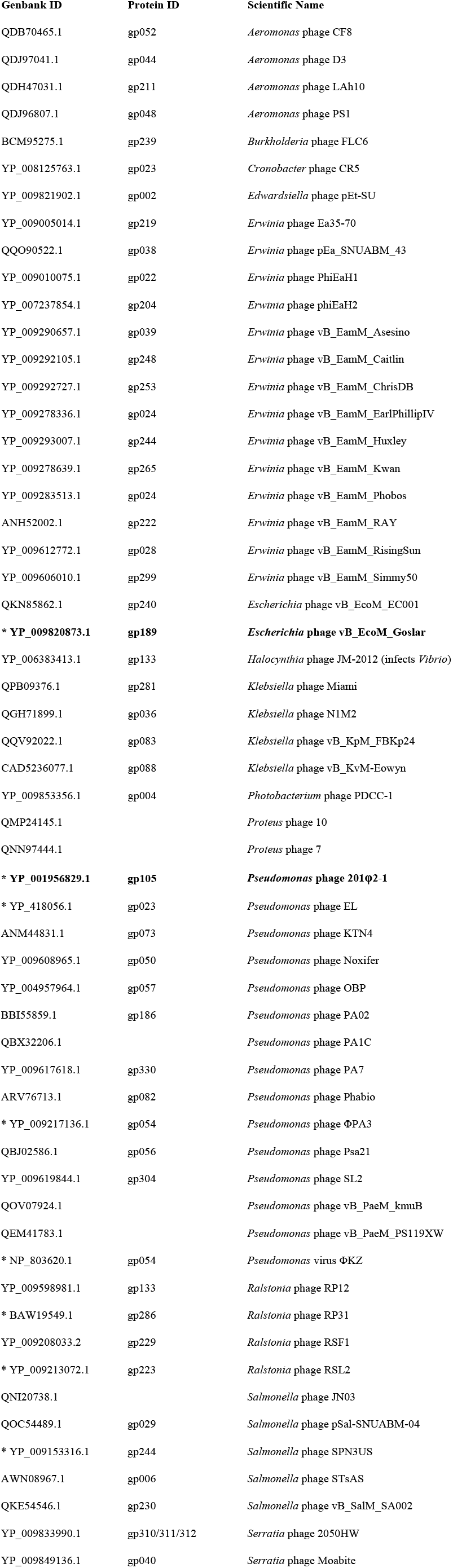
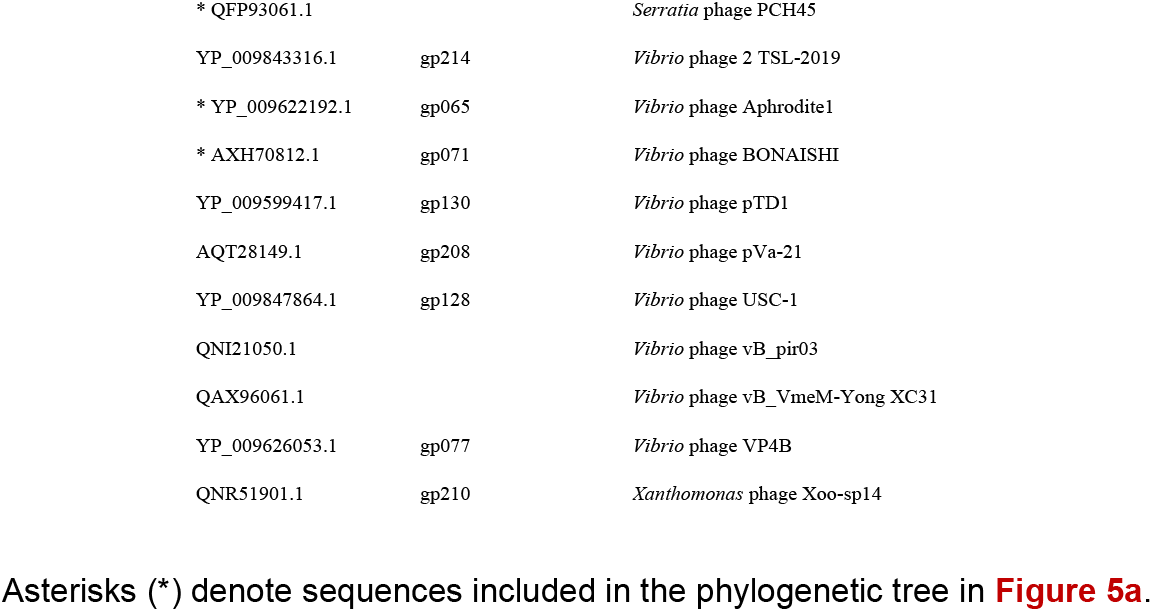
Bacteriophage chimallinA proteins.

| Genbank ID | Protein ID | Scientific Name |
| --- | --- | --- |
| QDB70465.1 | gp052 | *Aeromonas* phage CF8 |
| QDJ97041.1 | gp044 | *Aeromonas* phage D3 |
| QDH47031.1 | gp211 | *Aeromonas* phage LAh10 |
| QDJ96807.1 | gp048 | *Aeromonas* phage PS1 |
| BCM95275.1 | gp239 | *Burkholderia* phage FLC6 |
| YP_008125763.1 | gp023 | *Cronobacter* phage CR5 |
| YP_009821902.1 | gp002 | *Edwardsiella* phage pEt-SU |
| YP_009005014.1 | gp219 | *Erwinia* phage Ea35-70 |
| QQO90522.1 | gp038 | *Erwinia* phage pEa_SNUABM_43 |
| YP_009010075.1 | gp022 | *Erwinia* phage PhiEaH1 |
| YP_007237854.1 | gp204 | *Erwinia* phage phiEaH2 |
| YP_009290657.1 | gp039 | *Erwinia* phage vB_EamM_Asesino |
| YP_009292105.1 | gp248 | *Erwinia* phage vB_EamM_Caitlin |
| YP_009292727.1 | gp253 | *Erwinia* phage vB_EamM_ChrisDB |
| YP_009278336.1 | gp024 | *Erwinia* phage vB_EamM_EarlPhillipIV |
| YP_009293007.1 | gp244 | *Erwinia* phage vB_EamM_Huxley |
| YP_009278639.1 | gp265 | *Erwinia* phage vB_EamM_Kwan |
| YP_009283513.1 | gp024 | *Erwinia* phage vB_EamM_Phobos |
| ANH52002.1 | gp222 | *Erwinia* phage vB_EamM_RAY |
| YP_009612772.1 | gp028 | *Erwinia* phage vB_EamM_RisingSun |
| YP_009606010.1 | gp299 | *Erwinia* phage vB_EamM_Simmy50 |
| QKN85862.1 | gp240 | *Escherichia* phage vB_EcoM_EC001 |
| **\* YP_009820873.1** | **gp189** | ***Escherichia* phage vB_EcoM_Goslar** |
| YP_006383413.1 | gp133 | *Halocynthia* phage JM-2012 (infects *Vibrio*) |
| QPB09376.1 | gp281 | *Klebsiella* phage Miami |
| QGH71899.1 | gp036 | *Klebsiella* phage N1M2 |
| QQV92022.1 | gp083 | *Klebsiella* phage vB_KpM_FBKp24 |
| CAD5236077.1 | gp088 | *Klebsiella* phage vB_KvM-Eowyn |
| YP_009853356.1 | gp004 | *Photobacterium* phage PDCC-1 |
| QMP24145.1 |  | *Proteus* phage 10 |
| QNN97444.1 |  | *Proteus* phage 7 |
| **\* YP_001956829.1** | **gp105** | ***Pseudomonas* phage 201φ2-1** |
| * YP_418056.1 | gp023 | *Pseudomonas* phage EL |
| ANM44831.1 | gp073 | *Pseudomonas* phage KTN4 |
| YP_009608965.1 | gp050 | *Pseudomonas* phage Noxifer |
| YP_004957964.1 | gp057 | *Pseudomonas* phage OBP |
| BBI55859.1 | gp186 | *Pseudomonas* phage PA02 |
| QBX32206.1 |  | *Pseudomonas* phage PA1C |
| YP_009617618.1 | gp330 | *Pseudomonas* phage PA7 |
| ARV76713.1 | gp082 | *Pseudomonas* phage Phabio |
| * YP_009217136.1 | gp054 | *Pseudomonas* phage ΦPA3 |
| QBJ02586.1 | gp056 | *Pseudomonas* phage Psa21 |
| YP_009619844.1 | gp304 | *Pseudomonas* phage SL2 |
| QOV07924.1 |  | *Pseudomonas* phage vB_PaeM_kmuB |
| QEM41783.1 |  | *Pseudomonas* phage vB_PaeM_PS119XW |
| * NP_803620.1 | gp054 | *Pseudomonas* virus ΦKZ |
| YP_009598981.1 | gp133 | *Ralstonia* phage RP12 |
| * BAW19549.1 | gp286 | *Ralstonia* phage RP31 |
| YP_009208033.2 | gp229 | *Ralstonia* phage RSF1 |
| * YP_009213072.1 | gp223 | *Ralstonia* phage RSL2 |
| QNI20738.1 |  | *Salmonella* phage JN03 |
| QOC54489.1 | gp029 | *Salmonella* phage pSal-SNUABM-04 |
| * YP_009153316.1 | gp244 | *Salmonella* phage SPN3US |
| AWN08967.1 | gp006 | *Salmonella* phage STsAS |
| QKE54546.1 | gp230 | *Salmonella* phage vB_SalM_SA002 |
| YP_009833990.1 | gp310/311/312 | *Serratia* phage 2050HW |
| YP_009849136.1 | gp040 | *Serratia* phage Moabite |
| * QFP93061.1 |  | *Serratia* phage PCH45 |
| YP_009843316.1 | gp214 | *Vibrio* phage 2 TSL-2019 |
| * YP_009622192.1 | gp065 | *Vibrio* phage Aphrodite1 |
| * AXH70812.1 | gp071 | *Vibrio* phage BONAISHI |
| YP_009599417.1 | gp130 | *Vibrio* phage pTD1 |
| AQT28149.1 | gp208 | *Vibrio* phage pVa-21 |
| YP_009847864.1 | gp128 | *Vibrio* phage USC-1 |
| QNI21050.1 |  | *Vibrio* phage vB_pir03 |
| QAX96061.1 |  | *Vibrio* phage vB_VmeM-Yong XC31 |
| YP_009626053.1 | gp077 | *Vibrio* phage VP4B |
| QNR51901.1 | gp210 | *Xanthomonas* phage Xoo-sp14 |

Asterisks (\*) denote sequences included in the phylogenetic tree in **Figure 5a**.

**SI Table 10.** Bacterial strains and plasmids used in this study.

Asterisks (\*) denote sequences included in the phylogenetic tree in **Figure 5a**. SI Table 10 | Bacterial strains and plasmids used in this study.
| Bacterial strain | Source |
| --- | --- |
| *Pseudomonas chlororaphis* 200-B | (89) |
| *Escherichia coli* APEC 2248 | DSMZ (dsmz.de) #103255 |
| *Escherichia coli* Rosetta2 pLysS | EMD Millipore |

| Plasmid | Host strain | Source |
| --- | --- | --- |
| pHERD30-T | *P. chlororaphis* | (90) |
| pHERD30-T + GFPmut1 | *P. chlororaphis* | (91) |
| pHERD30-T + 201ϕ2-1 GFP-chimallinA full-length | *P. chlororaphis* | this study |
| pHERD30-T + 201ϕ2-1 GFP-chimallinA 48-631 | *P. chlororaphis* | this study |
| pHERD30-T + 201ϕ2-1 GFP-chimallinA 64-631 | *P. chlororaphis* | this study |
| pHERD30-T + 201ϕ2-1 GFP-chimallinA 1-611 | *P. chlororaphis* | this study |
| pHERD30-T + 201ϕ2-1 GFP-chimallinA 1-581 | *P. chlororaphis* | this study |
| pHERD30-T + 201ϕ2-1 GFP-chimallinA 48-611 | *P. chlororaphis* | this study |
| pHERD30-T + 201ϕ2-1 GFP-chimallinA 64-581 | *P. chlororaphis* | this study |
| 2-BT (His6-TEV site fusion expression vector) | *E. coli* Rosetta2 pLysS | Addgene (addgene.org) #29666 |
| 2-BT + 201ϕ2-1 chimallinA FL | *E. coli* Rosetta2 pLysS | this study |
| 2-BT + 201ϕ2-1 chimallinA 48-631 | *E. coli* Rosetta2 pLysS | this study |
| 2-BT + 201ϕ2-1 chimallinA 64-631 | *E. coli* Rosetta2 pLysS | this study |
| 2-BT + 201ϕ2-1 chimallinA 1-612 | *E. coli* Rosetta2 pLysS | this study |
| 2-BT + 201ϕ2-1 chimallinA 1-582 | *E. coli* Rosetta2 pLysS | this study |
| 2-BT + 201ϕ2-1 chimallinA 48-612 | *E. coli* Rosetta2 pLysS | this study |
| 2-BT + 201ϕ2-1 chimallinA 64-582 | *E. coli* Rosetta2 pLysS | this study |
| 2-BT + Goslar chimallinA full-length | *E. coli* Rosetta2 pLysS | this study |

## Video Legends

**Video 1** | **Sheet model anisotropic network model mode 1**

The sheet shows a scissoring motion, where the front of the sheet bends in one direction and the back of the sheet bends in the opposite direction. This movie used an RMSD difference of 25 Å from the original conformation.

**Video 2** | **Sheet model anisotropic network model mode 2**

The sheet arches downwards and upwards. This movie used an RMSD difference of 25 Å from the original conformation.

**Video 3** | **Top-down/cytosolic view of sheet model molecular dynamics trajectory**

Representative 300 ns molecular dynamics trajectory (n = 5) viewed from the top. Both the tetramers and their constituent monomers show a relative fluidity.

**Video 4** | **Tilted view of sheet model molecular dynamics trajectory**

Representative 300 ns molecular dynamics trajectory (n = 5) viewed from the side. The sheet shows wavelike undulations throughout the trajectory.

